# Multilevel community assembly of the tadpole gut microbiome

**DOI:** 10.1101/2020.07.05.188698

**Authors:** Decio T. Correa, David Rodriguez, Carine Emer, Daniel Saenz, Cory K. Adams, Luis Schiesari, Mikhail Matz, Mathew A. Leibold

## Abstract

The assembly of local communities is likely to reflect the effects of local environmental factors associated with filters that act at larger spatial scales. Dissecting these multiscale effects remains a timely challenge that is particularly important for host-associated microbiomes. We investigated the relative roles of local selection (due to host species identity) and regional effects (due to water body identity) on the community structure of bacteria in the gut of tadpoles from three biogeographic areas and used graph theory and metanetwork approaches to explore and illustrate the distribution of bacteria across different ponds. The pond of origin, which represents a regional species pool of bacteria, was in general more important in shaping the gut microbiome of tadpoles than host species identity. The resulting metanetworks are modular and indicate relatively few species of bacteria occurring in more than one pond. Thus, each pond represents a relatively distinct species pool of bacteria available for community assembly of the tadpole microbiomes. Our findings indicate that microbiome community assembly in amphibian larvae, as in many other communities, is a multiscale process with important regional effects that constrain how local (i.e. host-dependent) filters act to influence microbiome community composition.

## Introduction

The origin and assembly of host-associated microbial communities (microbiomes) is a complex process of fundamental importance in helping us understand the symbiotic relationship between hosts and their microbes [1, 2]. Various factors can play a role in the community assembly of microbiomes at multiple levels. Perhaps the most obvious is the effect of host diet on the gut microbiome, which has been shown in many organisms from insects to humans [3–6]. Other local factors include host sex, health state, and genetics [7–11]. These effects can be thought of as local ‘filters’ to microbial community assembly. Here we evaluate these effects as they are determined by host species identity only, recognizing that there are likely to be other more subtle effects involving inter-individual factors. However, there can also be processes that constrain which microbes are available to colonize any given host. Here, we focus on factors that determine the microbial species pool available to colonize the host and evaluate these by the identity of the site of the host community.

We view community assembly of microbiomes as being driven by metacommunity processes [12]. This recognizes that the assembly of a local community depends strongly on the species available to colonize that community, known as the regional species pool [13, 14]. From that pool, if a species is able to disperse to a given community, its persistence in that community will depend on local demographic processes and on the ability of the species to tolerate the local biotic and abiotic conditions, which is known as the local selection component of community assembly [15]. Therefore, communities with identical environmental conditions could have a completely different species composition if colonized by different species pools [16]. While most work in metacommunity ecology has focused on macroorganisms, these ideas should apply equally well to microbes [1, 2, 12, 17, 18].

Host-associated microbial communities appear to be influenced by effects related to the individual host or individual species such as diet, physiology, and genotype [10, 19, 20], as well as regional and historical processes, as in the case of exposure to potentially different species pools of microorganisms [21–26], or vertical and horizontal transmission of the microbiome [27, 28]. Considering the importance that the microbiome can have to host health and well-being, understanding the relative importance of effects acting at different levels is fundamental, but studies explicitly tackling that question are still lacking [29].

We analyzed variation in tadpole microbiomes to evaluate the effects of local environmental filters by focusing on host species identity, and regional effects by focusing on the locality of the water body. We assume that different microbes would be differentially favored by selection associated with the physiology and diet of different tadpole species, and different microbes may be present at different sites due to either site-specific habitat variables or to isolation and other differences among water bodies. Our studies were conducted across three different geographic locations to test if they show consistent patterns. We also used graph theory and metanetwork approaches to explore the degree to which each pond consists of a unique species pool, and how they are connected to each other in terms of shared bacterial species.

The gut microbiome of tadpoles is critically important to their digestive function [30], and some evidence shows gut microbiomes from the same host species are more similar to each other than from different species [31]. Further, there are often multiple species of tadpole per water body, which represent different hosts with different characteristics that are exposed to the same pool of bacteria. Many of these tadpole species can occupy a variety of different water bodies, each with potentially distinct bacterial species pools, which makes them a suitable system to test questions regarding the effects of environment and host identity on microbiome assembly [32]. We evaluate two hypotheses: i) if the bacterial community of each pond is different across the landscape, then the pond of origin would be more important in determining the gut microbiome of tadpoles, i.e., the microbiomes of different species within each pond would be more similar to each other than the microbiomes of the same species across distinct ponds, and the metanetworks would be structured into modules according to each pond; ii) if ponds have very similar bacterial communities across the landscape, then the species of tadpole would be more important in determining their gut microbiome, i.e., tadpoles of the same species across ponds would have more similar microbiomes than tadpoles of different species from the same pond, and the metanetworks would be structured into modules according to each species.

## Methods

### Study area

We sampled tadpoles from 22 water bodies located in three different localities: i) six water bodies in the Boracéia Biological Station, located in the Atlantic Forest east of the state of São Paulo, Brazil, hereafter BR; ii) nine water bodies in the David Crockett and Angelina National Forests, located in the pine forest ecosystem in eastern Texas, United States, hereafter ET; and iii) seven water bodies in the Edwards Plateau, a mix of grassland and juniper/oak woodlands in the central part of Texas, United States, hereafter CT. We classified water bodies as either lentic or lotic. For ET ponds, we did a more thorough environmental characterization quantifying in each pond the dissolved oxygen (mg/L), pH, conductivity (uS/cm), chlorophyll a (ug/L), and water temperature (°C) using a Eureka multiparameter sonde, and the absence or presence of piscivorous (Bass) and/or insectivorous fish (Green Sunfish). Even though three water bodies in BR were lotic, we will refer to all of them as *ponds* for convenience.

### Tadpole and microbiome sampling

In each pond we collected tadpoles with a dipnet that was swept through the pond. We then placed the larvae in a sterile plastic bag (Whirl Pak^®^) filled with water from their place of origin. Environmental microorganisms from water from CT and ET were sampled by filtering approximately 50 ml of water from three different places through a sterile syringe filter with a 0.45 μm cellulose acetate membrane (VWR). From BR, approximately 2L of water was collected in a sterile plastic bag (Whirl Pak^®^), mixed, and 100 mL was vacuum-filtered through a 0.45 μm cellulose acetate membrane. The sediment in all ponds was sampled in three locations in each pond by removing the first three centimeters with a sterile plastic straw of 1.3 cm of diameter and placed in a sterile tube.

In the lab, tadpoles were euthanized by overdose in a solution of Milli-Q^®^ filtered water and tricaine methanesulfonate (MS 222). After the death was confirmed, each individual was dissected using instruments that were cleaned in 100% ethanol and flame-sterilized. The entire tadpole gut was removed and placed directly in a MoBio PowerSoil^®^ bead tube. The sediment from each pond was homogenized, centrifuged to remove the excess water, and an aliquot of approximately 0.2 g was used for DNA extraction. DNA from the water sample was extracted from the entire filter membrane. We performed DNA extraction using MoBio PowerSoil^®^ DNA isolation kit following the manufacturer’s recommendation. The same procedures without a real sample were repeated to count as negative control samples, which did not have DNA when quantified using Qubit™dsDNA high sensitivity assay.

We amplified the V4 hypervariable region of the 16s rRNA gene (515F, 806F). CT samples were amplified using the version F: GTGYCAGCMGCCGCGGTA / R: GGACTACHVGGGTWTCTAAT and ET and BR the version F: GTGYCAGCMGCCGCGGTAA / R: GGACTACNVGGGTWTCTAAT. The library preparation and sequencing (Illumina MiSeq 2×250bp) of the CT samples was realized at the Genomic Sequencing and Analysis Facility (wikis.utexas.edu/display/GSAF) at The University of Texas at Austin following their protocols. The library preparation and sequencing (Illumina MiSeq 2×250bp) of the ET samples was realized at the Argonne National Laboratory (www.anl.gov) following their protocols. For BR samples, we prepared the libraries following the Earth Microbiome Project protocol (www.earthmicrobiome.org), with the exception of using 20uL PCR reactions and dual-index barcodes instead. Samples were sequenced in an Illumina MiSeq platform (2×300 bp paired-end sequences).

The sequences were processed using the dada2 pipeline [33] in the software Microsoft R Open v 3.5.1 [34]. Sequences from each site were processed separately. Briefly, primers, adapters, and barcodes were removed and the quality profile of the reads were visually inspected using the function plotQualityProfile aggregating over all the samples from a site. BR, CT, and ET reads were truncated at 200 and 120, 200 and 180, and 200 and 160 forward and reverse reads, respectively. Sequencing errors were calculated, sequences were clustered as Amplicon Sequence Variants based on the DADA2 algorithm [33] and paired ends were merged. Chimeras were removed using the consensus method in the function removeBimeraDenovo. Taxonomy was assigned based on the Silva database v128 [35]. ASV sequences were exported to Qiime2 2018.4 [36], aligned using the MAFFT algorithm [37], and a phylogenetic tree was constructed using FastTree [38]. We discarded all ASVs that were classified as Archaea, chloroplast, mitochondria, or that were not at least assigned to bacteria. Hereafter, we will refer to ASVs as species of bacteria.

## Data analysis

### Diversity

For each sample we constructed a rarefaction curve and compared richness and Faith’s phylogenetic diversity [39]. To avoid sequencing depth effects, each sample was rarefied 100 times to the number of reads of the sample with the lowest coverage using the R package BAT [40]. For comparison, we considered the median richness from all rarefactions from a sample as our richness measure.

### Species and pond effects

We analyzed our data as compositional data (termed CODA) [41]. It requires a centered log-ratio transformation of the read counts and therefore cannot have zeros, so we added a pseudocount of one to all ASVs in all samples. We transformed the data using the codaSeq.clr function from the R package CoDaSeq [42, 43]. We also analyzed data that takes into account phylogenetic relatedness of microbes by using the PHILR transformation from the R package philr [44]. This metric takes into account the phylogenetic relationship between the ASVs, which is equivalent to the Unifrac metric [45], but it considers the compositional nature of the data [44]. We then ran a Principal Component Analysis (PCA) on the Euclidean distance matrix of the CODA- and PHILR-transformed data to visualize the relationship between samples and tested if samples from tadpoles, water, and sediment are different from each other using a Permutational Multivariate Analysis of Variance (perMANOVA). If a difference was detected we then used additional perMANOVAs for pairwise comparisons with Bonferroni correction.

To first test if either the identity of the species of tadpole or the pond of origin are more related to the gut microbiome composition we used a perMANOVA to fit a model with both pond and species identity to the CODA- and PHILR-transformed data using the function adonis2 from the R package vegan [46]. The significance of the marginal effects was tested based on 999 permutations. In addition, to test for the unique effects of the species of tadpole or the pond of origin as well their shared effects on the gut microbiome we performed variance partitioning and Redundancy Analysis (RDA) [47] using the function varpart in the R package vegan [46]. The significance of each unique component in the variation partitioning was tested using the vegan function anova.cca with 1000 permutations. We repeated perMANOVA and variation partitioning analyses to the data grouping the species of bacteria at Genus and Family level.

### Inter-pond variation

We also investigated if there was evidence for dispersal limitation in the bacteria from water and sediment using multiple regression on distance matrices using the function MRM from the R package ecodist [48]. We used the Euclidean distance matrix based on the PHILR and CODA transformations as the dependent variable and a geographical distance matrix between sites as the independent variable. In addition, to account for environmental effects, we also include in the model a distance matrix based on the type of the pond (lentic or lotic) for BR, and a distance matrix based on the environmental variables measured for ET. The latter was a Gower distance matrix [49] created by the function dist.ktab from the R package ade4 [50]. Quantitative variables were Gower-standardized (divided by maximum minus minimum). All CT ponds were lentic and there were no other variables besides spatial distance to include in the model. For models where distance was significant in the MRMs, we applied a Partial Mantel Correlogram approach using the function mpmcorrelogram from the R package with the same name [51]. That tests, at several distance classes, in our case estimated by Sturge’s rule [52], the relationship between water and sediment microbiome with geographical distance while controlling for environmental effects.

### Metanetworks

We further explored the structure of three different metanetworks within each locality (BR, CT, and ET) to understand the distribution of bacteria across ponds. Shared species (here bacteria) connect different ponds in a metacommunity perspective [53, 54]. We built three bacteria metanetworks for each locality: (i) the tadpole microbiome metanetwork, composed by the bacteria living in the gut of all tadpoles within a pond, independently of the tadpole species, (ii) the water metanetwork, with bacteria found in the water, and (iii) the sediment metanetwork, composed by bacteria found in the sediment of each pond. Then, we built an a_*mn*_ adjacency matrix for each metanetwork in which *m* corresponds to a single pond (i.e., considering bacteria from water, sediment or gut of tadpoles), and *n* to each bacterium. The *mn* element represents the presence/absence of species of bacteria *n* in a pond *m*, and is represented by a link in the graphical representation. Therefore, each node in our metanetwork is either a bacterium or a pond; when the same bacterium is found in more than one pond, there is a link connecting them, forming the edges of the metanetwork.

To test whether ponds consist of different bacterial communities, we calculated the modularity of the metanetworks using the FastGreedy algorithm [55, 56] in the software Modular v 0.1 [57]. We tested the significance of the observed modularity against two different null models with 100 permutations each: the Erdős-Rényi [58], in which pond-bacteria interactions are connected randomly, and the ‘null model 2’ [59], in which the probability of a pond-bacteria interaction is proportional to their number of links in the observed matrix. To identify the bacteria with higher potential to connect the metanetwork through shared occurrence in a higher number of ponds, we calculated the degree (k) for each metanetwork using the software Pajek v. 4.10 [60]. Degree measures how many links each node establishes in its correspondent network. For the bacteria, it represents the number of ponds in which it occurs; for the pond, it represents the number of bacteria it has (from tadpoles, water or sediment). For each metanetwork we also calculated the proportion of bacteria found uniquely in p_1_, p_2_, …, p_n_, where n is the total number of ponds in one locality.

## Results

### Diversity

We registered 19 species of tadpole in all ponds: three in CT, six in ET, and eleven in BR (Table S1). After the filtering steps, there were a total of 533 777, 1 289 662, and 2 064 298 16s rRNA gene fragment reads for CT, ET, and BR, respectively. The rarefaction curve shows that all samples reached an asymptote, even the ones with the lowest number of reads (Figure S1). For richness and phylogenetic diversity measures of bacteria, CT, ET, and BR samples were rarefied to 3 950, 1 791, and 3 916 reads, respectively. There was no significant difference among CT samples in terms of richness, but there was in terms of phylogenetic diversity, with sediment samples having the highest diversity and *Rana berlandieri* tadpoles the lowest (Figure S2). Sediment samples had the highest richness and phylogenetic diversity in ET and *Hylodes phyllodes* the highest in BR (Figures S3 and S4). A great part of the tadpole microbiome was composed by Fusobacteria, Firmicutes, Proteobacteria, and Bacteroidetes, while sediment and water samples were mostly composed by Proteobacteria (Figures S5-S7).

### Species and pond effects

The composition of the tadpole gut microbiome is different from both water and sediment samples in all localities (Figure 1, Figure S8, Table S2). Also, in most cases, tadpole gut microbiome samples cluster more with other samples from the same pond than with samples from the same species from a different pond (Figure 1, Figures S8-S10). However, within a pond, there can be separation by species and, in fact, the perMANOVA shows that both species identity of tadpole and pond of origin are significant predictors of the tadpole gut microbiome in all localities when using either PHILR- or CODA-transformed data, except for CT CODA, in which species identity was only marginally significant (Table 1). In CT and ET, pond of origin was a stronger predictor of the tadpole gut microbiome than tadpole species identity. The overall results are consistent over all the taxonomic levels considered (Tables S3-S6).

**Table 1.** Results of a Permutational Analysis of Variance (perMANOVA) testing if the species of tadpole or the pond of origin are related to the gut microbiome composition of tadpoles using the bacteria as Amplicon Sequence Variants (ASVs). Models were fitted to CODA and PHILR transformed data. The significance of the marginal effects was tested based on 999 permutations. BR = Brazil, CT = Central Texas, ET = East Texas.

|  |  | PHILR |  |  | CODA |  |  |
| --- | --- | --- | --- | --- | --- | --- | --- |
|  |  | R<sup>2</sup> | F | p | R<sup>2</sup> | F | p |
| BR | Water body<sub>(4,80)</sub> | 0.08 | 3 | 0.005 | 0.1 | 2.7 | 0.001 |
|  | Species<sub>(9,80)</sub> | 0.15 | 2.5 | 0.002 | 0.16 | 2.1 | 0.001 |
| CT | Water body<sub>(5,25)</sub> | 0.48 | 6 | 0.001 | 0.41 | 3.6 | 0.001 |
|  | Species<sub>(1,25)</sub> | 0.05 | 2.8 | 0.010 | 0.04 | 1.8 | 0.056 |
| ET | Water body<sub>(8,76)</sub> | 0.27 | 4.7 | 0.001 | 0.24 | 3.2 | 0.001 |
|  | Species<sub>(5,76)</sub> | 0.18 | 5.1 | 0.001 | 0.12 | 2.6 | 0.001 |

**Figure 1.**
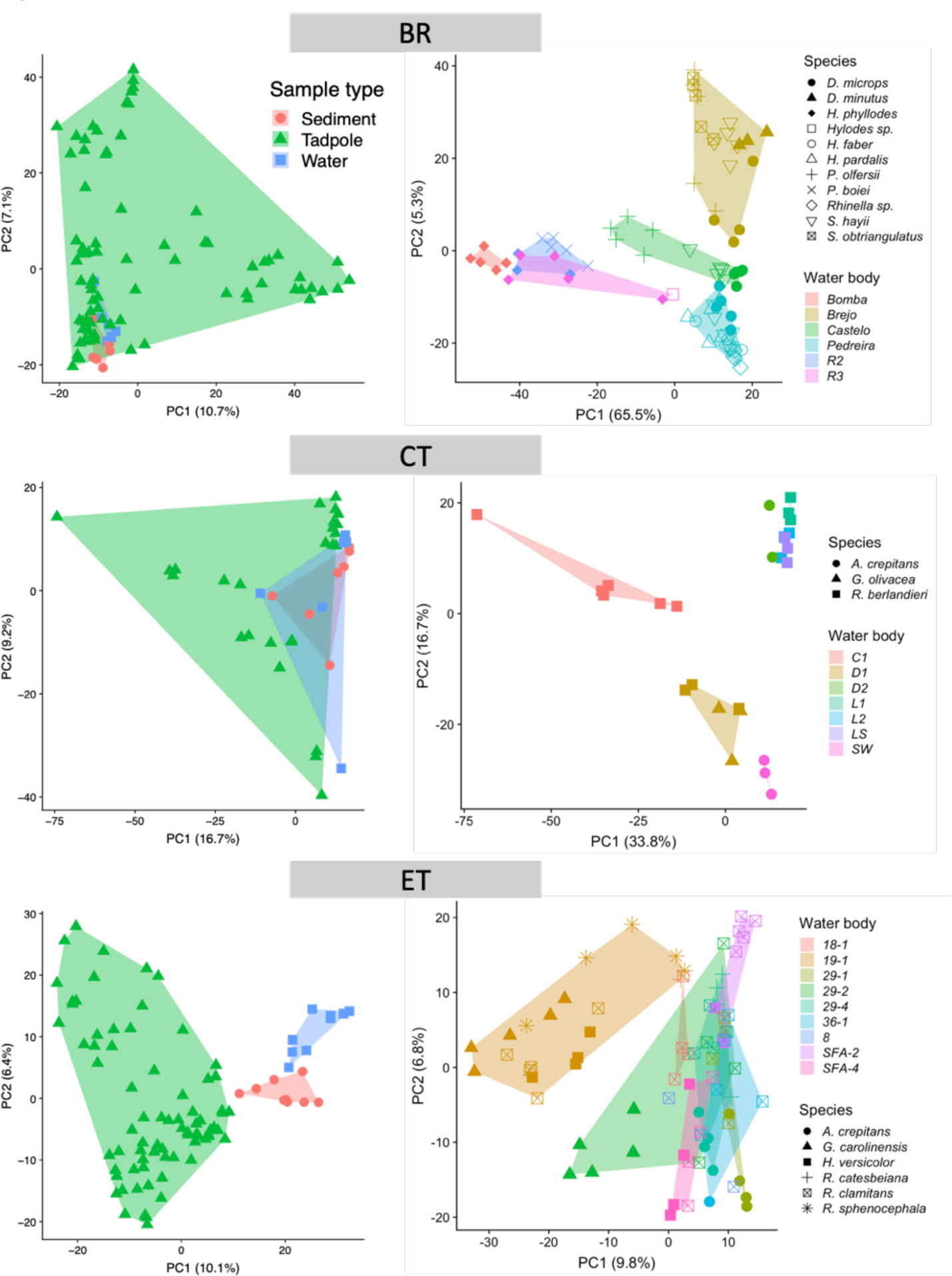
Plot of the first two axes of a Principal Component Analysis on CODA transformed microbiome data (amplicon sequence variants) from tadpole samples from Brazil (BR), Central Texas (CT), and Eastern Texas (ET). Panels on the left show sediment, water, and tadpole samples. Panels on the right show samples from tadpoles only. Left and right panels are results from separate analyses. Shapes represent different species of tadpoles and colors represent water body of origin. See Table S1 for abbreviations.

For CT and ET, the pond of origin explained considerably more of the variance in tadpole gut microbiome than the species of tadpole (Figure 2). For BR, the tadpole species explained a slightly higher proportion of the variance in their gut microbiome. However, most of the variation in that environment is explained by the joint effects of tadpole species and pond of origin, i.e., the variation that could not be separated in unique components (Figure 2). Again, the overall results are comparable across all the taxonomic levels considered (Tables S7-S8).

**Figure 2.**
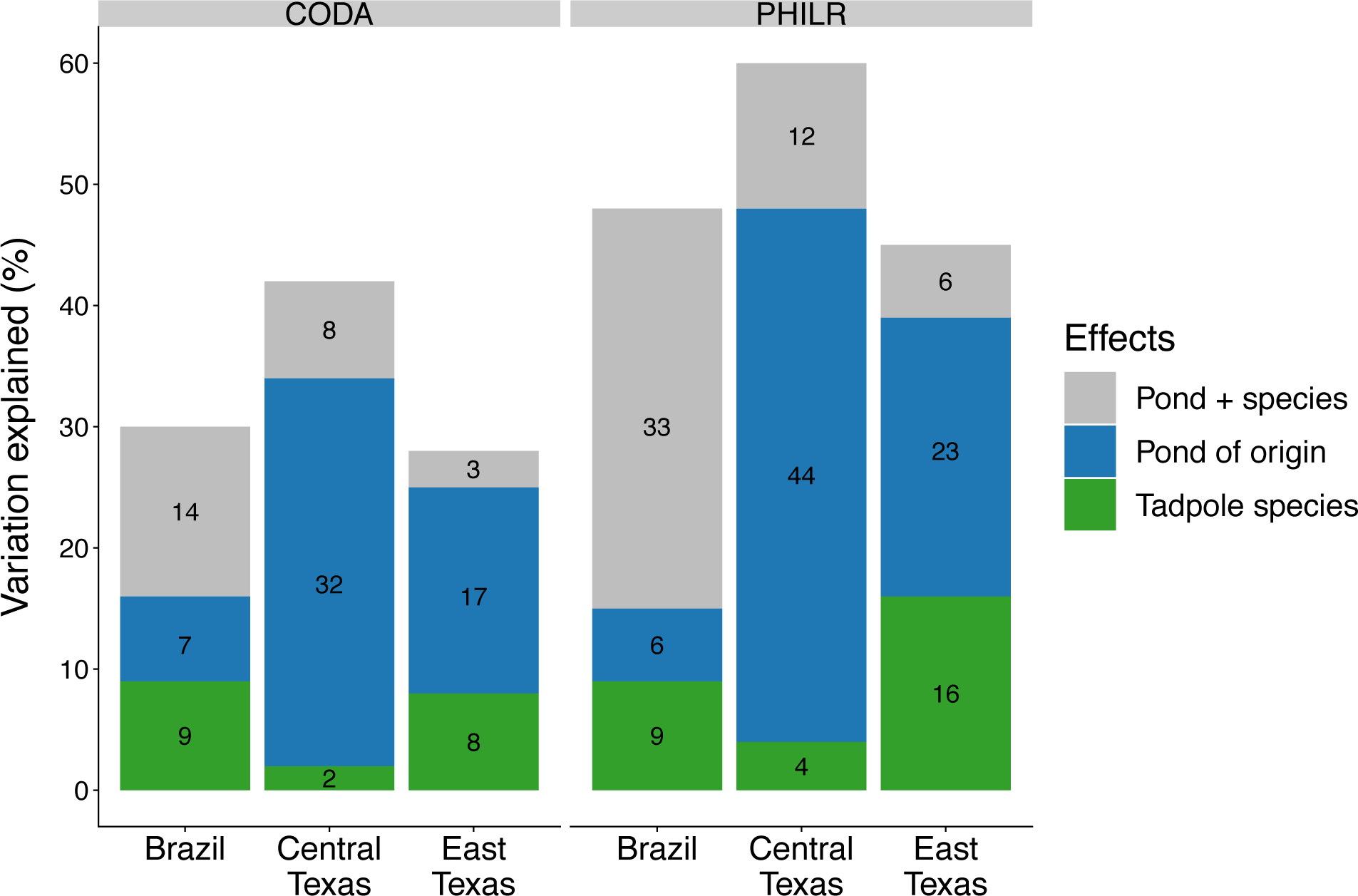
Results of a variation partitioning analysis testing the unique effects of the species of tadpole or pond of origin as well their shared effects on the tadpole gut microbiome considering the bacteria as Amplicon Sequence Variants (ASVs) for CODA- and PHILR-transformed data. The effects of pond of origin and species of tadpole were all significant (p < 0.05) based on 1000 permutations, except for the shared component, which cannot be tested.

### Inter-pond variation

Only the bacterial communities of water from CT were significantly related to the spatial distance between ponds (Table S9). However, that was the only dataset where environmental data is not available and could not be accounted for, allowing for the possibility that such distance effects are due to spatially structured environmental factors. Partial Mantel Correlograms showed that, for both CODA- (p_adj_ = 0.044) and PHILR-transformed data (p_adj_ = 0.022), there is a spatial signal only for the first distance range, from 174 to 6 223 m. ET water and sediment communities were significantly related to environmental variables for PHILR-transformed data and BR water communities were significantly related to both PHILR- and CODA-transformed data (Table S9).

### Metanetworks

All metanetworks are significantly modular according to the two null models used (p < 0.001 for all null models; Figure 3, Figure S11). In most cases, each module corresponds to a single pond (Figure 3, Figure S11 and Table S10). Metanetworks of water and sediment are more modular than those of tadpoles for all localities (Figure 3, Figure S11). In all cases, the vast majority of bacteria occur in a single pond (*k* = 1), varying from 63.2% in East Texas to 91.3% in Brazil (Figure 4, Figures S12-S16). Only 0.1 to 1.1% of bacteria were present in all ponds (Figure 4, Figure S12-S16). From those, for BR and ET they were mostly from the phyla Bacteroidetes and Proteobacteria and Orders Bacteroidales and Desulfovibrionales. CT had only three bacteria present across all ponds. From the ponds’ perspective, their degree, which corresponds to the number of unique links they have in the metanetwork, varied from 118 to 615 for water, 90 to 478 for sediment, and 241 to 2 751 for bacteria from tadpoles (Figures S17-S19).

**Figure 3.**
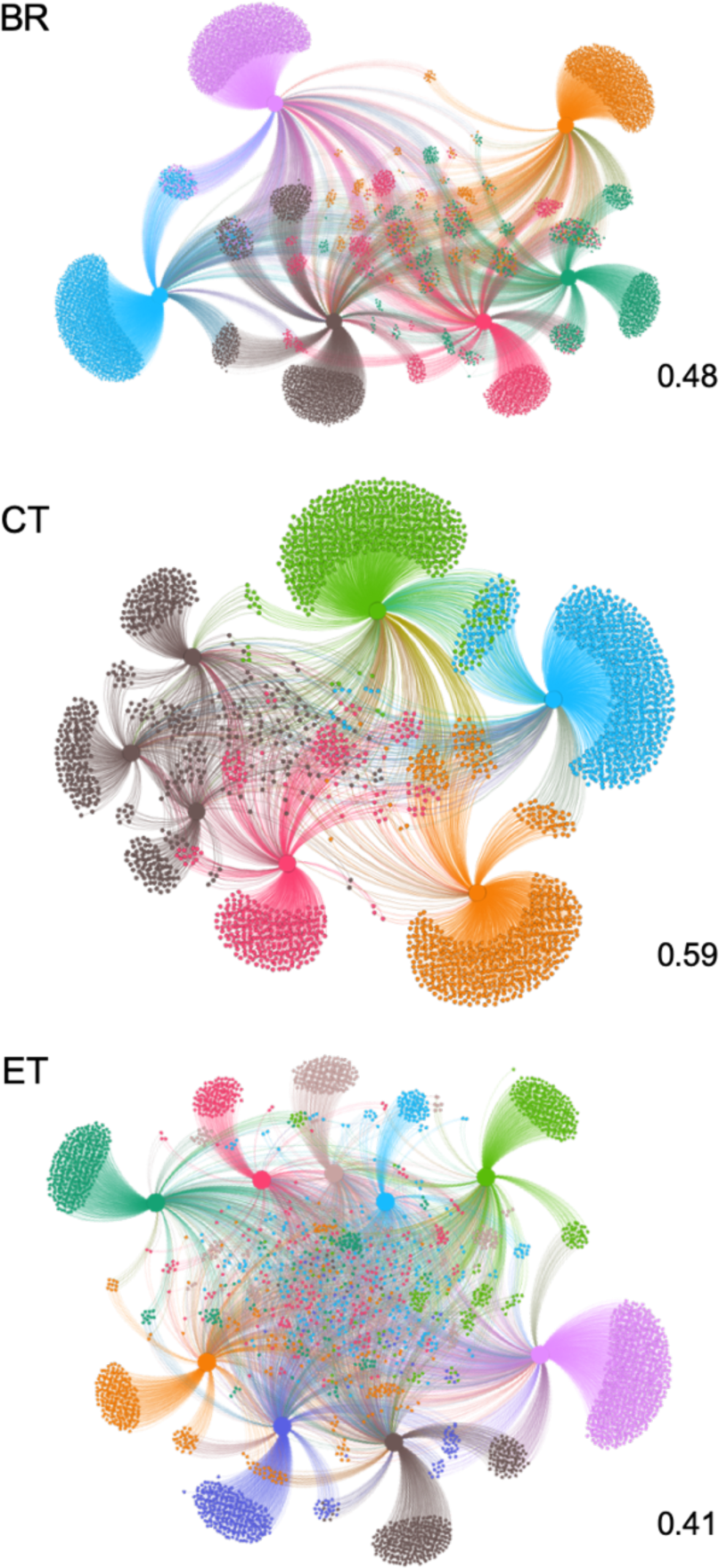
Microbiome metanetworks of the shared bacteria (amplicon sequence variant - ASV) found in the gut of tadpoles. Larger nodes represent each water body for each locality, *i*.*e*., Brazil (BR), Central Texas (CT), and East Texas (ET). Smaller nodes represent a unique bacterium that can be found in a single water body (link between smaller and larger nodes) or shared among ponds thus connecting them. Numbers on the bottom-right are the modularity values for each metanetwork. Distinct colors represent different modules which, with a few exceptions, correspond to a single water body (Table S10).

**Figure 4.**
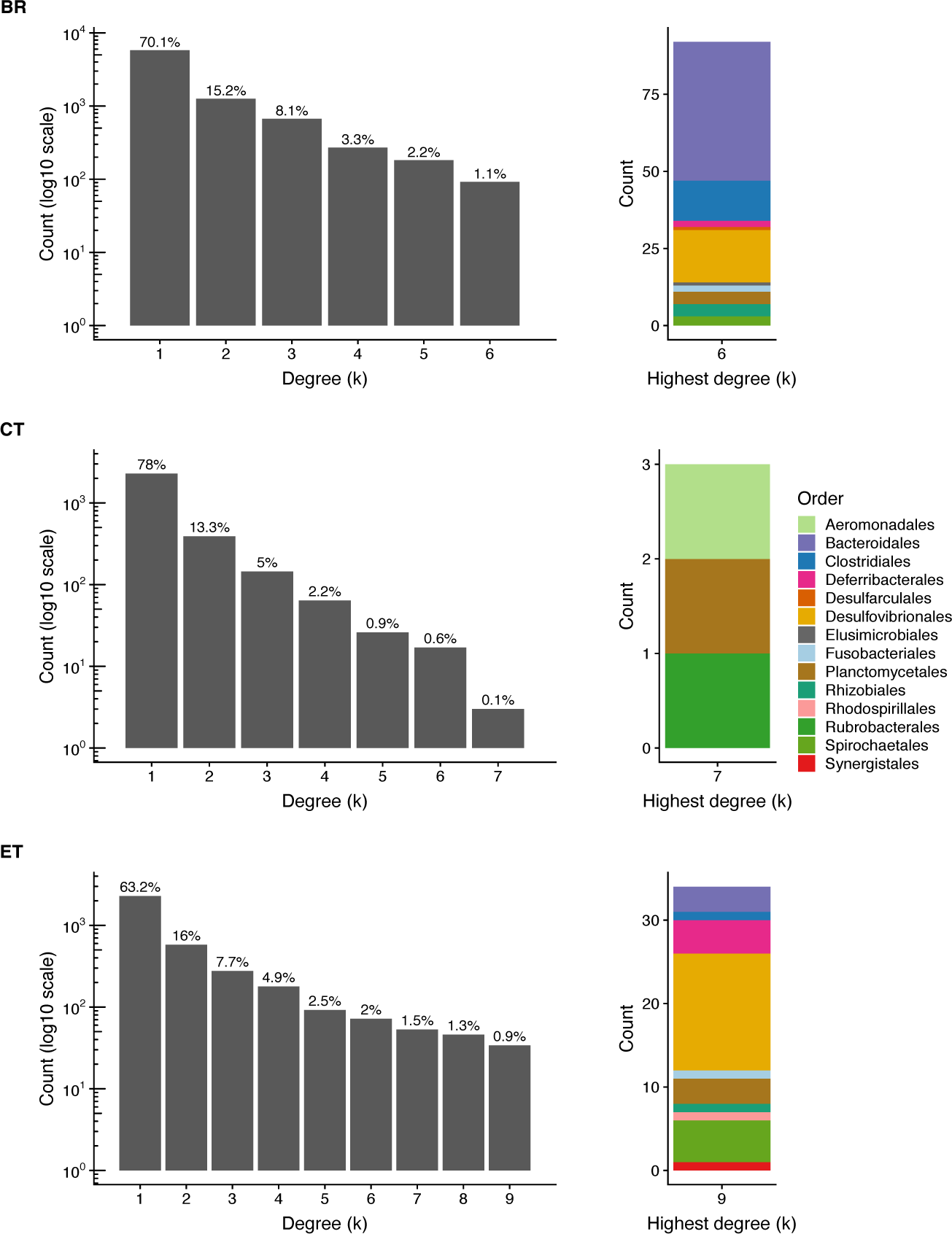
Number of bacteria from the gut of tadpoles for each degree (k) on the left and Order of bacteria present in the highest degree on the right. Each degree represents one pond. Bacteria found in only one pond are represented in the degree category 1. Likewise, bacteria present in tadpoles across all ponds are represented in the highest category of each histogram. The Order of bacteria present in tadpoles across all ponds from each locality is shown on the barplot on the right. Same color across barplots represent the same Order. See Figures S11-S16 for water and sediment metanetworks. BR = Brazil, CT = Central Texas, ET = East Texas.

## Discussion

We found that microbial community assembly of tadpole microbiomes was structured by processes happening at different levels and broadly resemble findings from macroorganisms [1, 15, 61–63]. At higher levels, there are inter-pond differences in microbial communities due to site-specific variables, isolation, and/or stochasticity. It was not our goal to understand the drivers of inter-pond differences, which is ultimately responsible for the different bacterial species pool tadpoles are exposed to. However, in some cases, for the ponds where we had environmental data there is a relationship between environmental variables and the bacterial community of water and/or substrate. In these cases, each pond could act as a regional scale environmental filter for bacteria. For CT, we found some evidence for a spatial correlation in bacterial communities among nearby ponds, which could indicate dispersal limitation structuring the pond communities, but that was the only dataset for which we did not have environmental variables. Nevertheless, independently of the causes of the differences among ponds, each one of them constitutes a unique species pool of colonizers for the microbiomes of tadpoles. As tadpoles develop, the community assembly of their microbiome is restricted by that species pool of microbes. Each tadpole comprises a smaller-scale filter to a subset of the microbes within the species pool. Therefore, even if similar habitats (same species of tadpole in our case) are exposed to different species pools, they will not end up with the same microbial community.

The composition and diversity of the gut microbiome of tadpoles can be variable across species. The microbiome of tadpoles from this study is slightly different than previous studies. Proteobacteria and Firmicutes dominated in other studies [31, 64–66] but here we found that Fusobacteria were also important. Alpha and phylogenetic diversity were also mostly higher in our study than in previous ones [31, 64–66]. The knowledge about the gut microbiome of tadpoles is still limited to a few species and quantitative comparisons of diversity across studies is problematic because of differences in sampling procedures, sample processing, data cleaning, and the algorithms and databases. However, overall differences in microbiome composition could be due to the exposure of tadpoles to different pools of colonist microbes.

The pond of origin of the tadpoles was more important in determining their gut microbiome than the species of the tadpole in both of the Texas locations. We assume the same species of tadpole has similar requirements in terms of microbes and/or constitutes a similar patch for colonization by microbes. There were only a few bacteria in common across all ponds in all three localities, therefore each pond constitutes a unique bacterial species pool from which the tadpoles can be colonized. In Brazil, the pond of origin and species of tadpole showed similar relationships with the gut microbiome of tadpoles. However, most of variation in Brazil was due to variation that could not be separately attributed to either because of the uniquely distinct tadpole faunas of ponds.

The limited and unique pool of bacteria tadpoles are exposed to and the higher similarity within a pond can be visualized on the metanetworks of bacteria from water, sediment, and tadpole gut. The metanetworks are modular and each pond constitutes a unique community of bacteria. The vast majority of bacteria occur only in a single pond, and just a few are present in all of them. In fact, for water and sediment, the maximum number of bacteria found in all ponds was only eight. That number would probably increase if more sediment or water was sampled. On the other hand, we likely sampled all species of tadpoles from each water body and the proportion of bacteria from their gut found in all water bodies is still very low (0.1-1.1%) despite similar phyla found in all of them. The major bacteria in tadpole guts belong to orders Bacteroidales and Desulfovibrionales, which do not overlap with the most widespread taxa found in water or sediment. It is common to not find much overlap between bacteria from environment and gut of aquatic organisms [67, 68]. Likewise, we show that the gut microbiome of tadpoles is distinct from both water and sediment from the ponds suggesting that the gut constitutes a strong environmental filter that favors certain taxa. Desulfovibrionales, for example, is a group of sulfur-reducing bacteria that is mostly anaerobic [69]. Therefore, at least certain parts of the digestive tract of tadpoles might be anaerobic [31, 70]. Pryor and Bjorndal [30] detected significant levels of fermentation in the hindgut of bullfrog tadpoles. It is unknown to which extent other species of tadpoles are hindgut fermenters. For those that are, even though they might not have a core microbiome, at least not at fine-scale taxonomy levels, there may be a functional core gut microbiome.

In terrestrial and aquatic environments, the importance of the regional species pool in shaping the microbial communities can be noticed even when not tested directly. In many species, from insects to humans, individuals likely exposed to the same regional species pool, such as being from the same environment, have more similar microbiomes [21–26]. There are many difficulties in quantifying and testing how regional effects affect community assembly [29, 62], but comparing results with predictions from theory can lead to important insights. If hosts from different environments have similar microbiomes it could indicate that (i) there is not much difference in the regional species pools between environments, (ii) the hosts move between species pools, or (iii) there are other methods for microbiome acquisition such as transmission from conspecifics. For example, Kueneman *et al*. [71] and McKenzie *et al*. [32] studying the skin microbiome of tadpoles found that the identity of the species was the strongest predictor of the skin microbiome, with the water body of origin explaining additional [71] or no variance at all [32]. Vertical transmission of the microbiome is unlikely in amphibians [72], and they probably acquire their microbiome from the environment at every generation [73]. Therefore, the results from Kueneman *et al*. [71] and McKenzie *et al*. [32] indicate there is at least a partially shared species pool of microbes across their sampling sites.

The enormous intra and inter-specific variation in the microbiomes of different species of animals and plants can be at least partially explained by differences in regional species pool during the community assembly of the microbiome. The gut microbiome of animals can be stable over time [63, 74] and can also be resistant to invasions and colonization by other bacteria [75–77]. Therefore, if the community assembly of individuals of the same species occurred while they were exposed to different species pools, historical effects can cause lasting effects in the development of the microbiome [78]. Moreover, recent efforts to manipulate the microbiome of plants and animals to improve health and/or increase productivity might not be as effective if the manipulation is done only at the level of the individual host, i.e., at the local scale. As stated by Chase [16] “if local and regional processes determine the community composition, then both processes need to be restored to achieve the desired community”. Our results indicate that the same logic is valid for microbes.

In summary, we showed that local assembly of host-associated bacterial communities are affected by regional scale processes, more specifically changes in the regional species pool of colonizers. The inter-pond variation in bacterial community can be due to stochastic, historical, spatial, and/or environmental processes or even the result of local evolution [79]. Nevertheless, microbiome assembly, as any other community, is a multiscale process. Microbial and microbiome ecology could be better linked to ecological theory by considering the multiscale dynamics of community assembly [12, 17, 80, 81]. Because it is much easier to quantify effects at the scale of the individual host or host species, i.e., local effects [29, 62], the role of regional effects is probably underestimated. Regional effects, however, appear to be a fundamental piece of the community assembly puzzle that can be used to understand variation in microbiomes.

## Acknowledgements

We thank Ananda Brito de Assis and Carlos Arturo Navas for laboratory access during field work in Brazil and Christopher Schalk and James Childress for help in the field in East Texas. We thank the Zoology Museum of University of São Paulo (MZUSP), The National Forests and Grasslands of Texas, and several private landowners for granting permission to access sampling sites and ICMBio (47263-1), Texas Parks and Wildlife (SPR-0414-067), and IACUC (AUP-2014-00286) for collecting and animal care permits. Funding was provided by Conselho Nacional de Desenvolvimento Científico e Tecnológico (CNPq, Brasil, grant number 458796/2014-0) and Texas Ecolab. DTC received a Science Without Borders scholarship during the research (CAPES 1176/13-7). Finally, we thank Daniel Bolnick, Michel Ryan, Ulrich Mueller, and members of Leibold and Matz labs for comments throughout the project.

