## Supplemental material for "Multilevel community assembly of the tadpole gut microbiome"

### Supplementary information

### Figures

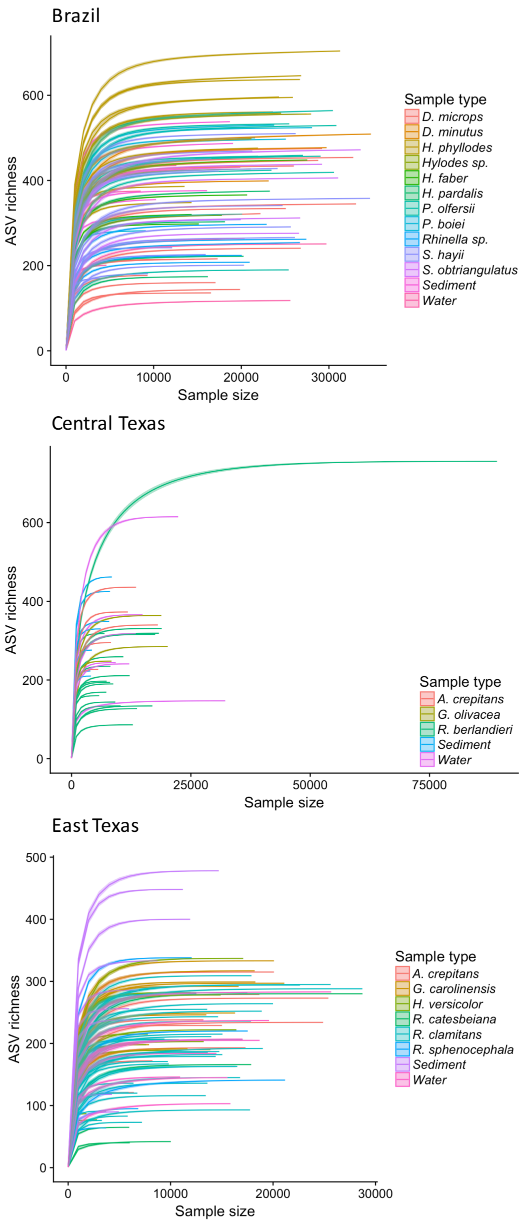


Figure S1. Rarefaction curves for each locality for all the Amplicon Sequence Variants classified at least as bacteria (see Methods). Shaded region on the curves represent standard error. Sample size is measured in number of reads.

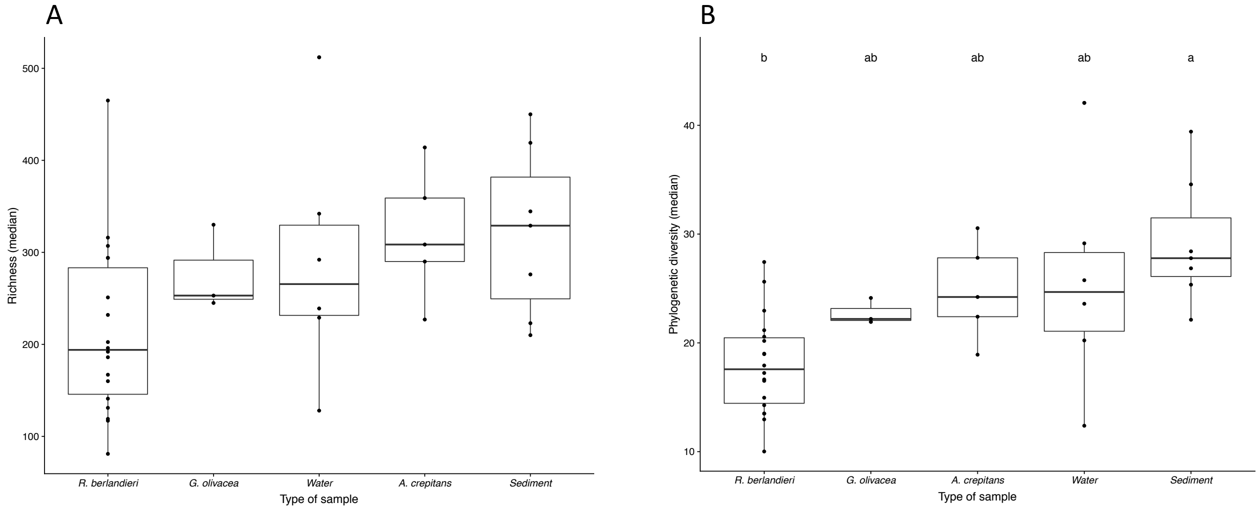


Figure S2. Boxplot of median values of amplicon sequence variant richness (panel A) and Faith's phylogenetic diversity (panel B) per type of sample from Central Texas. Dots are raw values. Different letters on top of each boxplot represent significant difference between each type of sample (Tukey HSD post-hoc test, alpha = 0.05). See Table S1 for abbreviations.

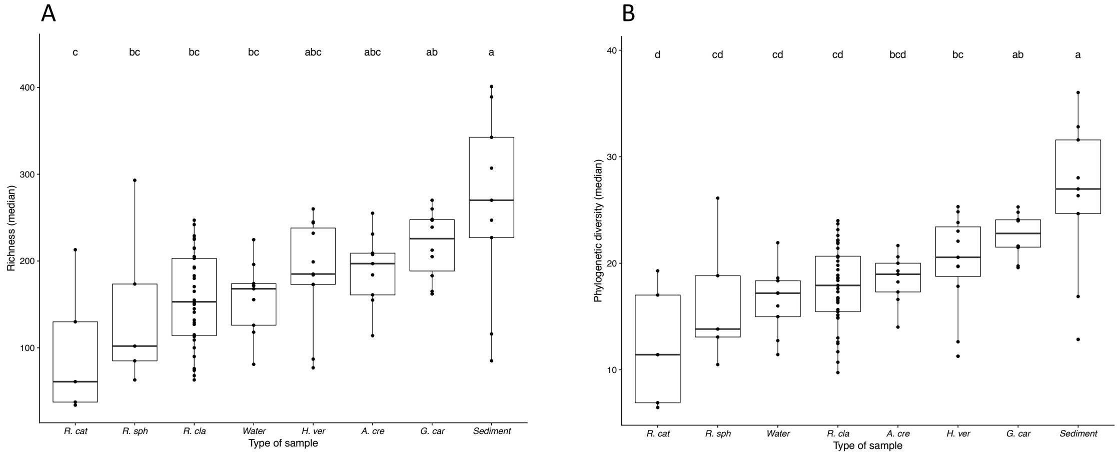


Figure S3. Boxplot of median values of amplicon sequence variant richness (panel A) and Faith's phylogenetic diversity (panel B) per type of sample from East Texas. Dots are raw values. Different letters on top of each boxplot represent significant difference between each type of sample (Tukey HSD post-hoc test, alpha = 0.05). See Table S1 for abbreviations.

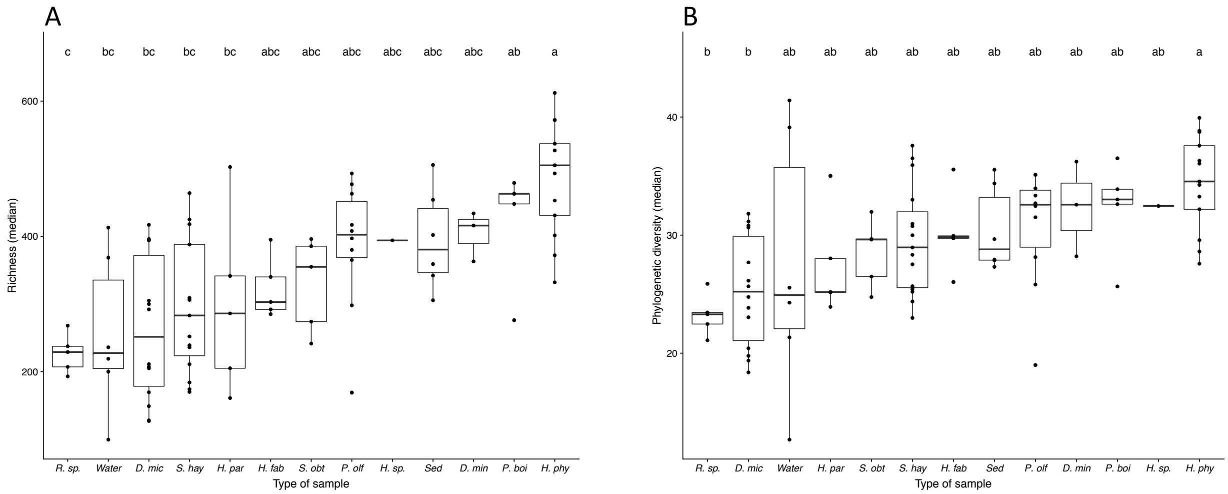


Figure S4. Boxplot of median values of amplicon sequence variant richness (panel A) and Faith's phylogenetic diversity (panel B) per type of sample from Brazil. Dots are raw values. Different letters on top of each boxplot represent significant difference between each type of sample (Tukey HSD post-hoc test, alpha = 0.05). See Table S1 for abbreviations.

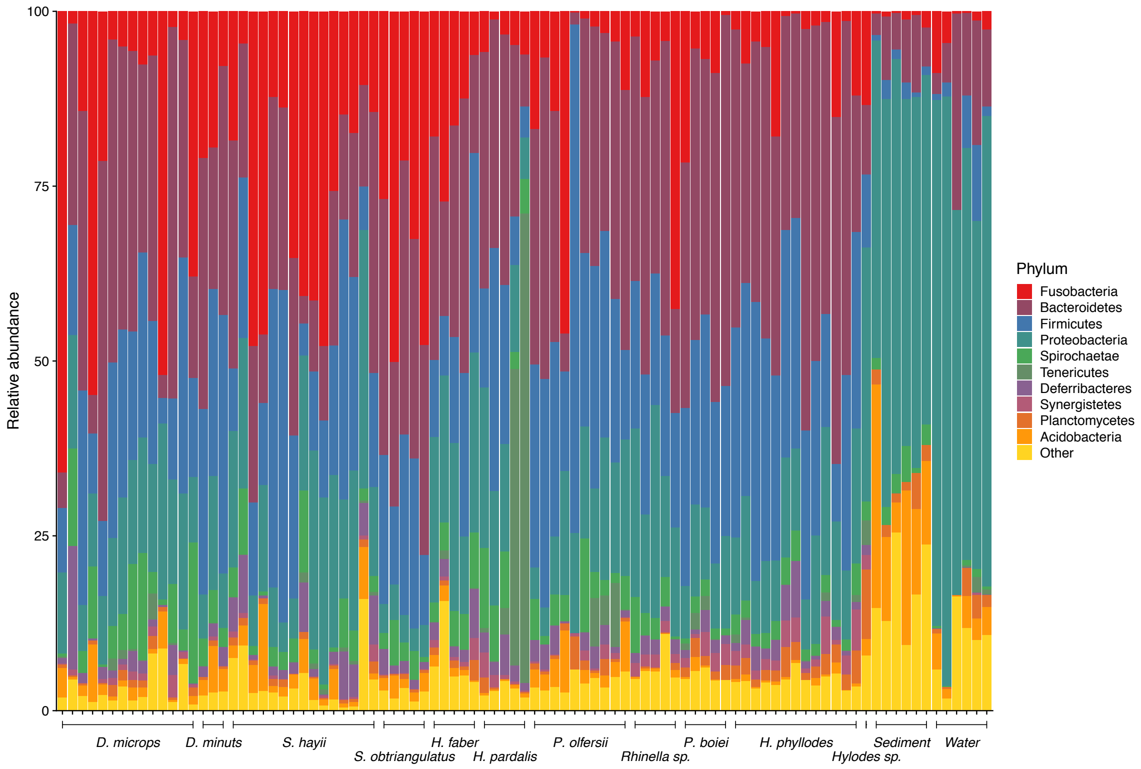


Figure S5. Relative abundance of the main phyla of bacteria found in each sample from Brazil. See Table S1 for abbreviations.

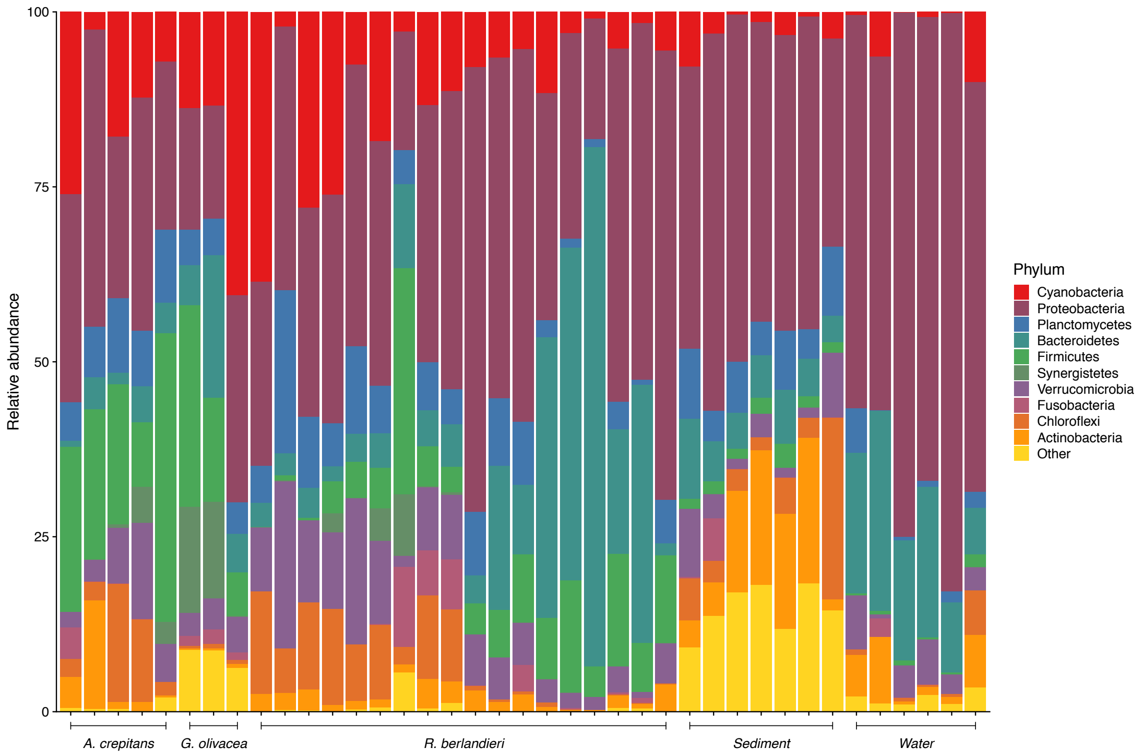


Figure S6. Relative abundance of the main phyla of bacteria found in each sample from Central Texas. See Table S1 for abbreviations.

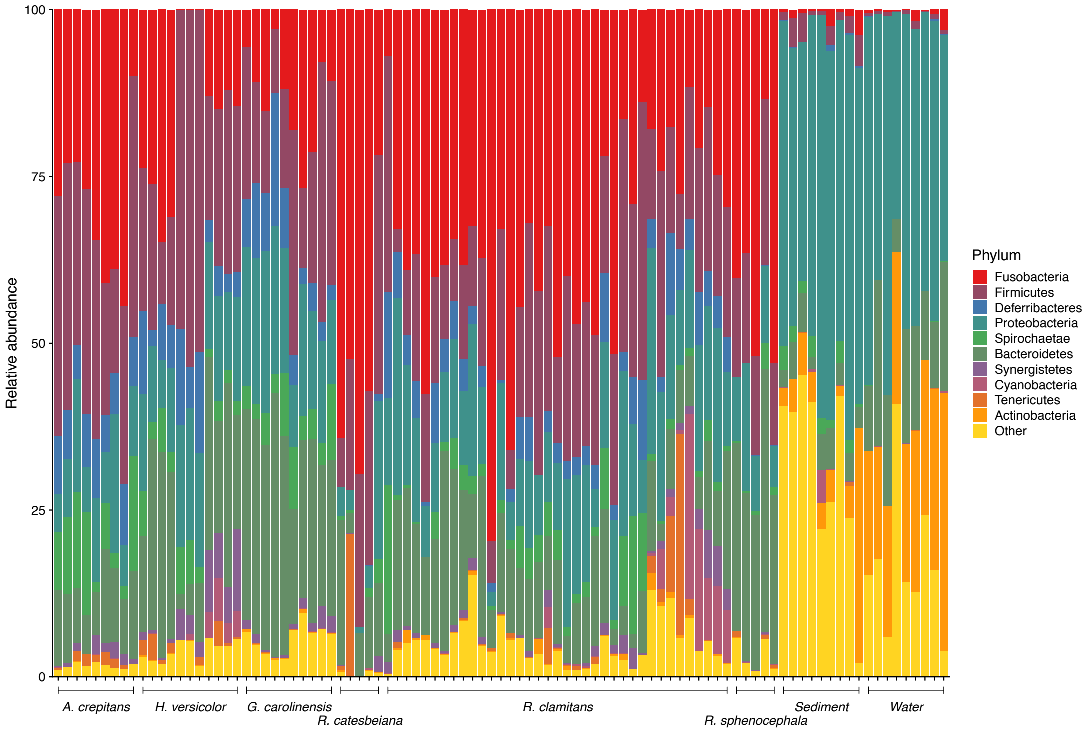


Figure S7. Relative abundance of the main phyla of bacteria found in each sample from East Texas. See Table S1 for abbreviations.

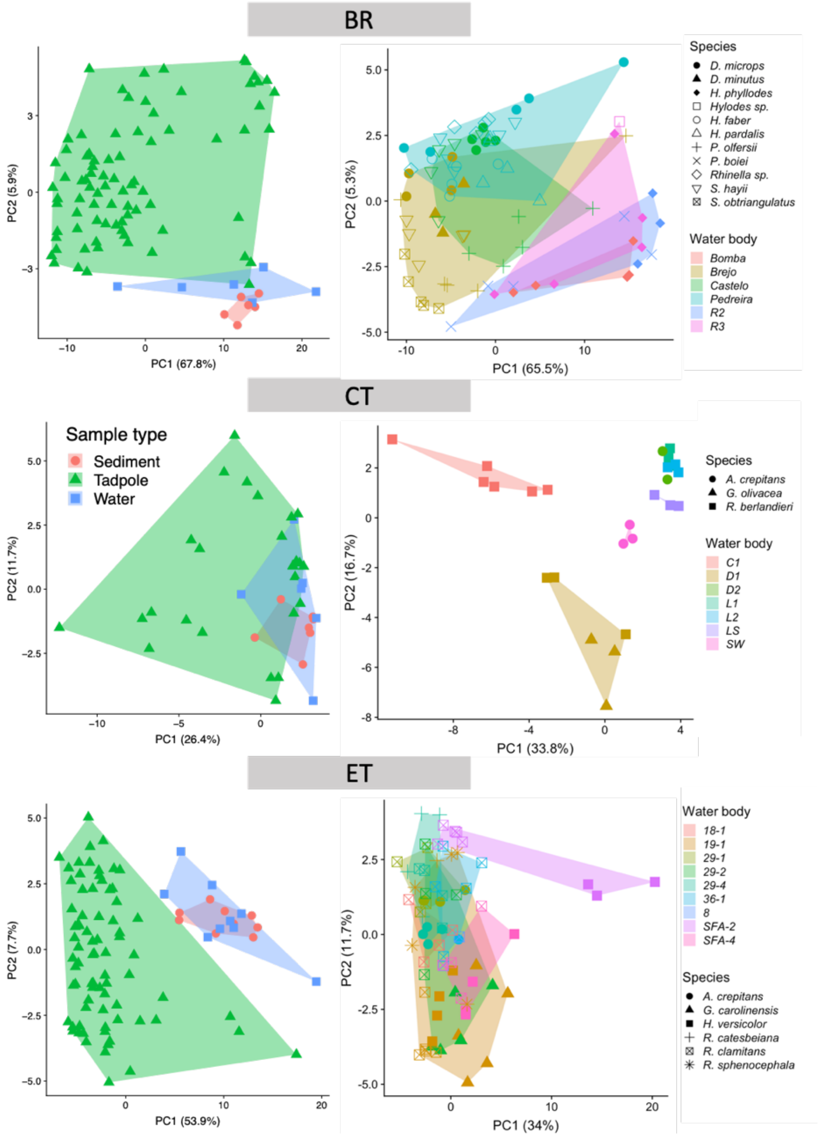


Figure S8. Plot of the first two axes of a Principal Component Analysis on PHILR transformed microbiome data (amplicon sequence variants) from tadpole samples from Brazil (BR), Central Texas (CT), and Eastern Texas (ET). Panels on the left show sediment, water, and tadpole samples represented by different shapes and colors. Panels on the right show samples from tadpoles only. Shapes represent different species of tadpoles and colors represent water body of origin. See Table S1 for abbreviations

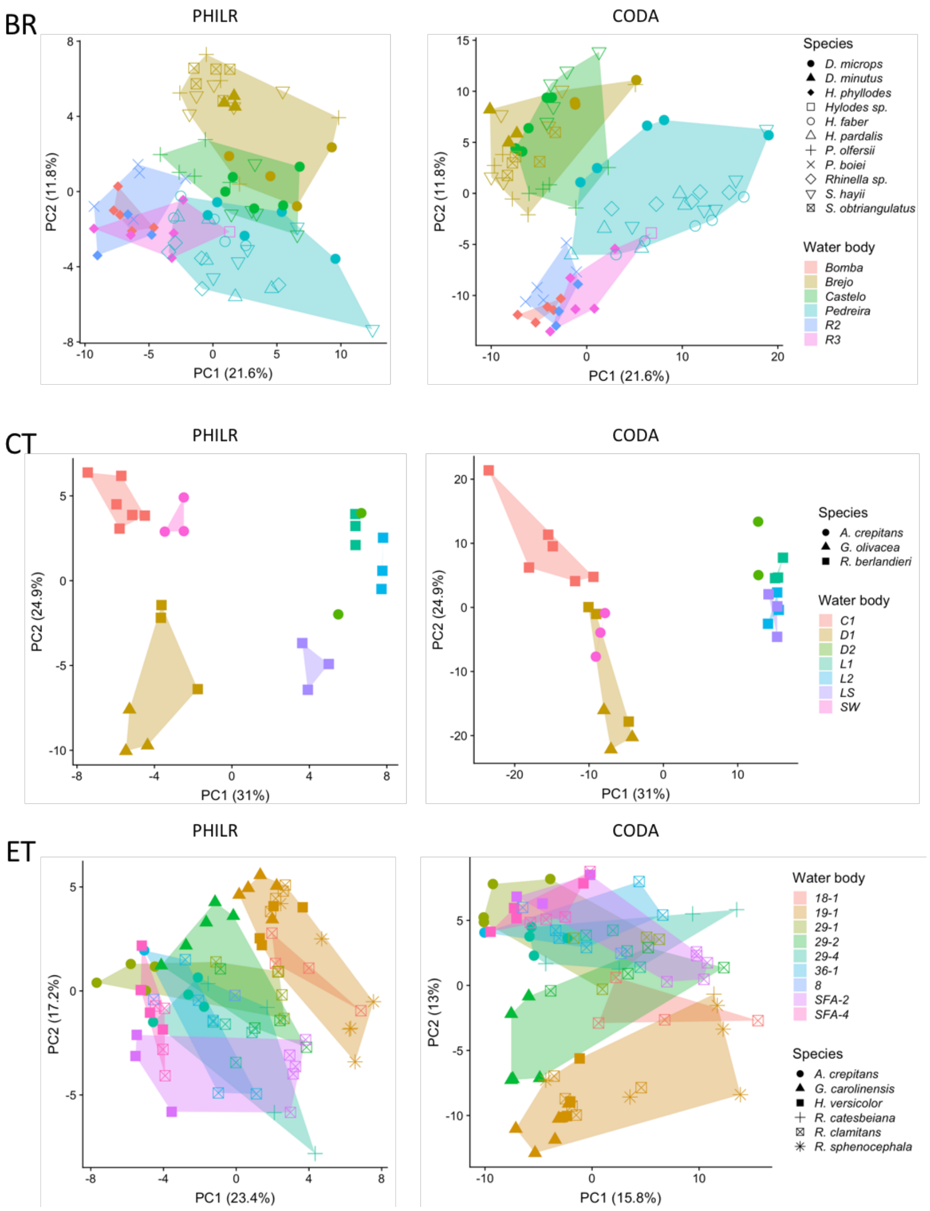


Figure S9. Plot of the first two axes of a Principal Component Analysis on PHILR and CODA transformed microbiome data (amplicon sequence variants grouped by Genus) from tadpole samples from Brazil (BR), Central Texas (CT), and Eastern Texas (ET). Shapes represent different species of tadpoles and colors represent water body of origin. See Table S1 for abbreviations.

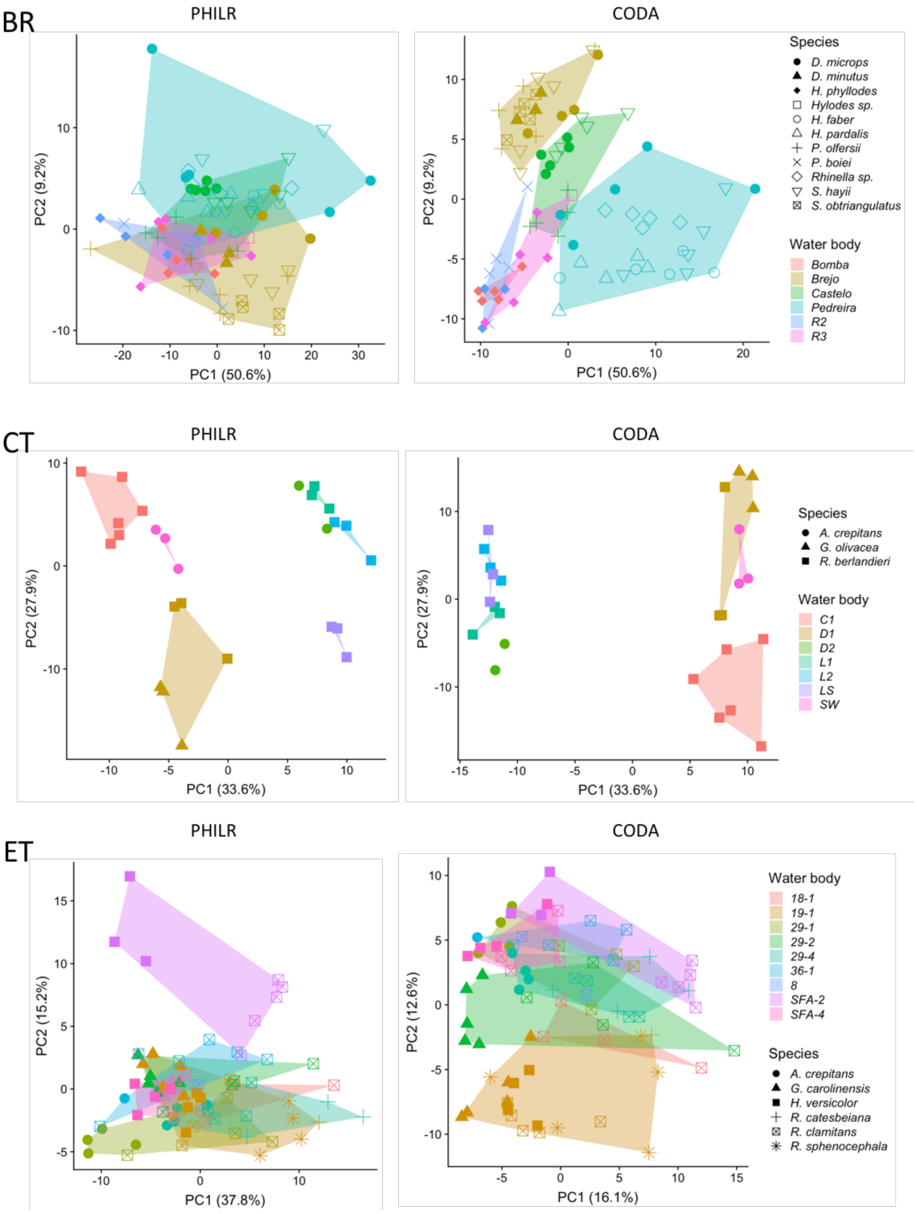


Figure S10. Plot of the first two axes of a Principal Component Analysis on PHILR and CODA transformed microbiome data (amplicon sequence variants grouped by Family) from tadpole samples from Brazil (BR), Central Texas (CT), and Eastern Texas (ET). Shapes represent different species of tadpoles and colors represent water body of origin. See Table S1 for abbreviations.

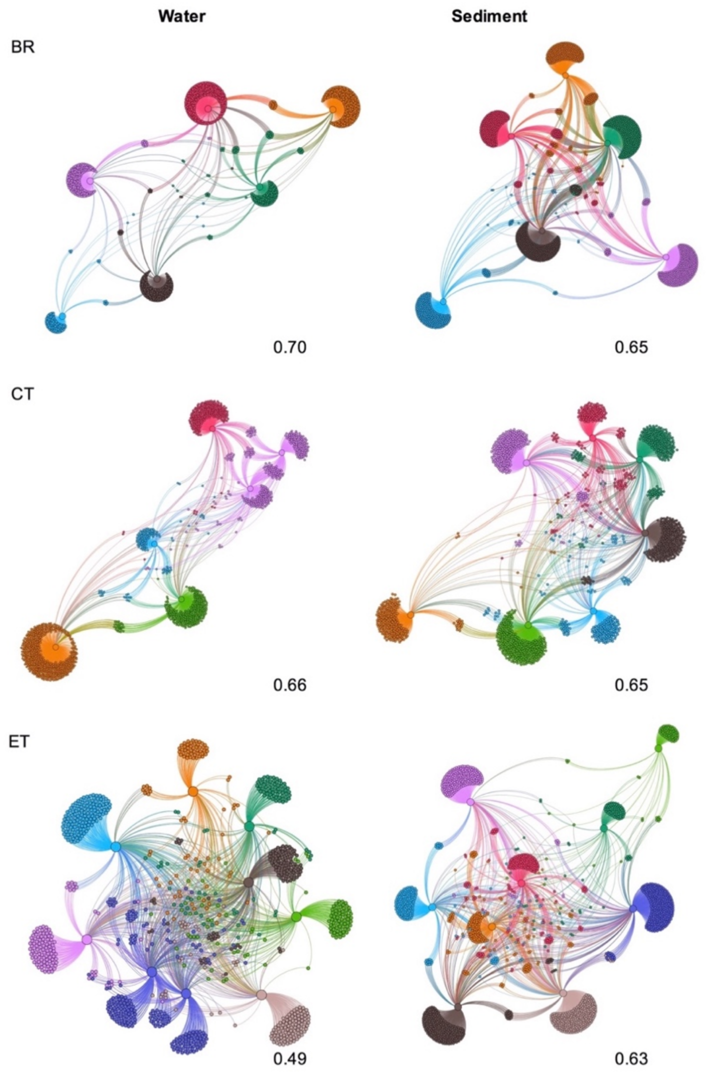


Figure S11. Graph representation of the water and sediment networks. Large circles represent each water body. Networks are significantly modular (P < 0.001) and each module is represented by a different color, which, with a few exceptions, correspond to a single water body (Table S10). Smaller circles represent a bacterium (ASV) that is shared across water bodies if there is a line connecting them. Number on the bottom right of networks are the modularity values. BR = Brazil, CT = Central Texas, ET = East Texas.

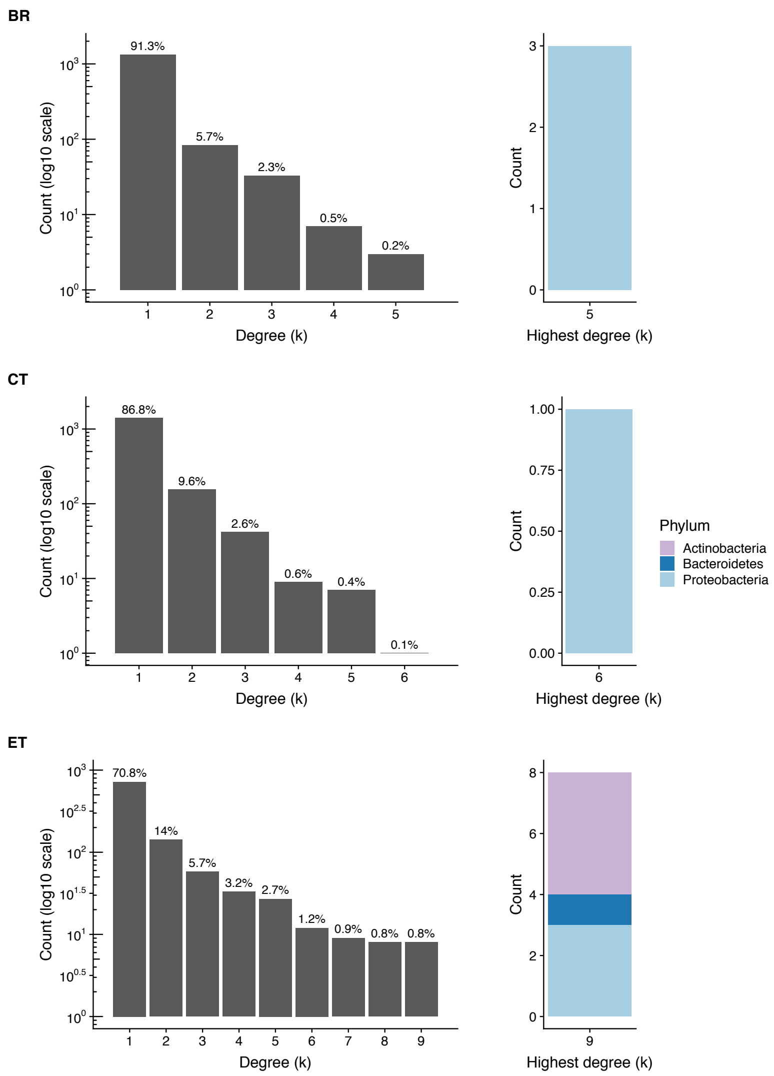


Figure S12. Number of ASVs from water for each degree (k) on the left and Phylum of ASVs present in the highest degree on the right. Each degree represents one water body. ASVs found in only one water body are represented in the category 1. Likewise, ASVs present in water across all water bodies are represented in the highest category of each histogram. Note that the count for each histogram is in log10 scale. The Phylum of each ASV present in water across all water bodies from each locality is shown on the barplot on the right. Same color across barplots represent the same Phylum. BR = Brazil, CT = Central Texas, ET = East Texas.

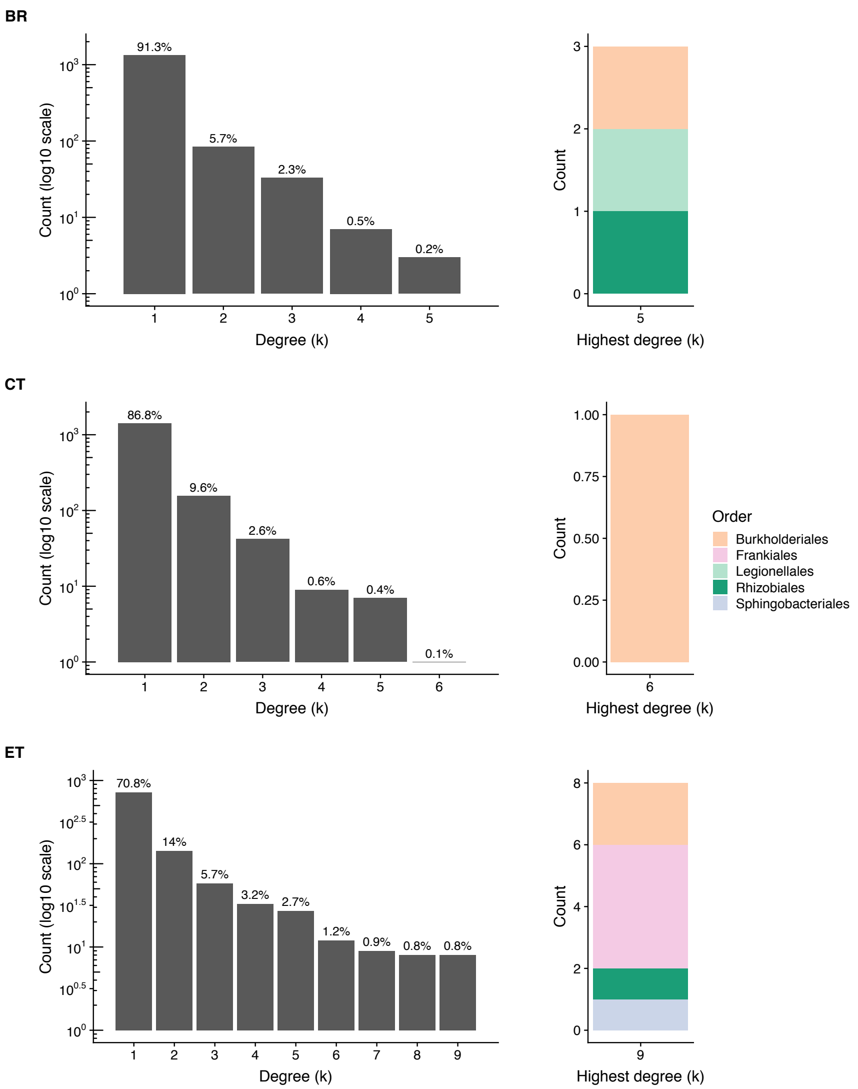


Figure S13. Number of ASVs from water for each degree (k) on the left and Order of ASVs present in the highest degree on the right. Each degree represents one water body. ASVs found in only one water body are represented in the category 1. Likewise, ASVs present in water across all water bodies are represented in the highest category of each histogram. Note that the count for each histogram is in log10 scale. The Order of each ASV present in water across all water bodies from each locality is shown on the barplot on the right. Same color across barplots represent the same Order. BR = Brazil, CT = Central Texas, ET = East Texas.

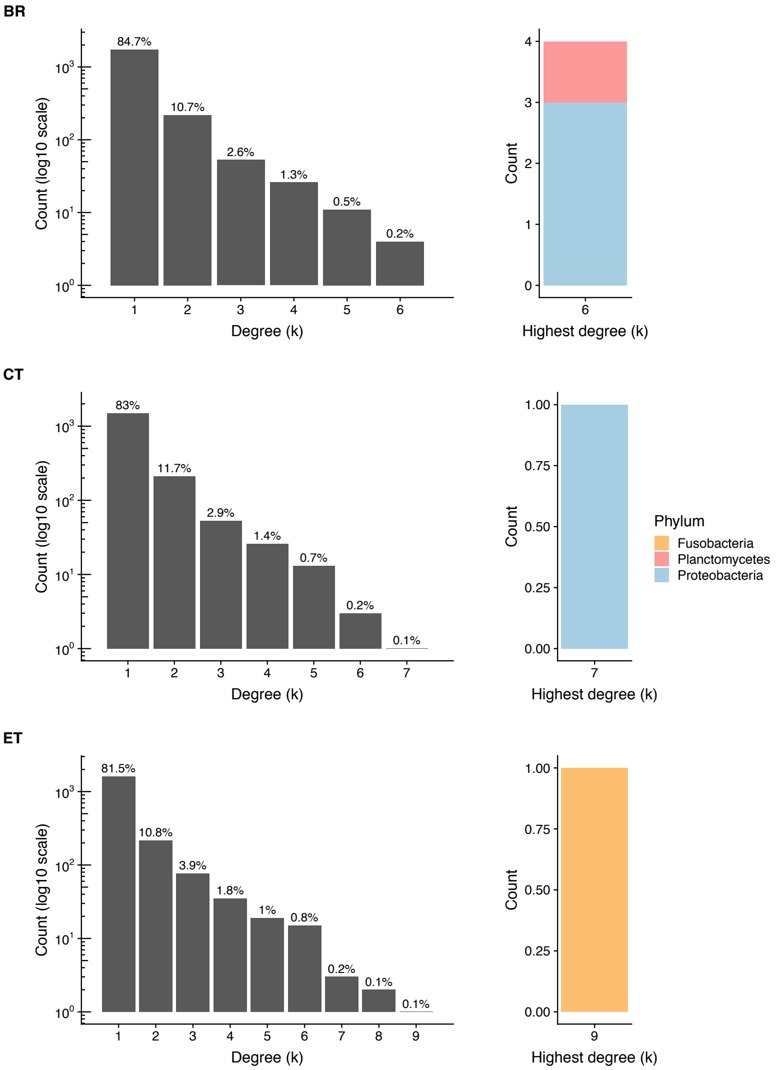


Figure S14. Number of ASVs from sediment for each degree (k) on the left and Phylum of ASVs present in the highest degree on the right. Each degree represents one water body. ASVs found in only one water body are represented in the category 1. Likewise, ASVs present in sediment across all water bodies are represented in the highest category of each histogram. Note that the count for each histogram is in log10 scale. The Phylum of each ASV present in sediment across all water bodies from each locality is shown on the barplot on the right. Same color across barplots represent the same Phylum. BR = Brazil, CT = Central Texas, ET = East Texas.

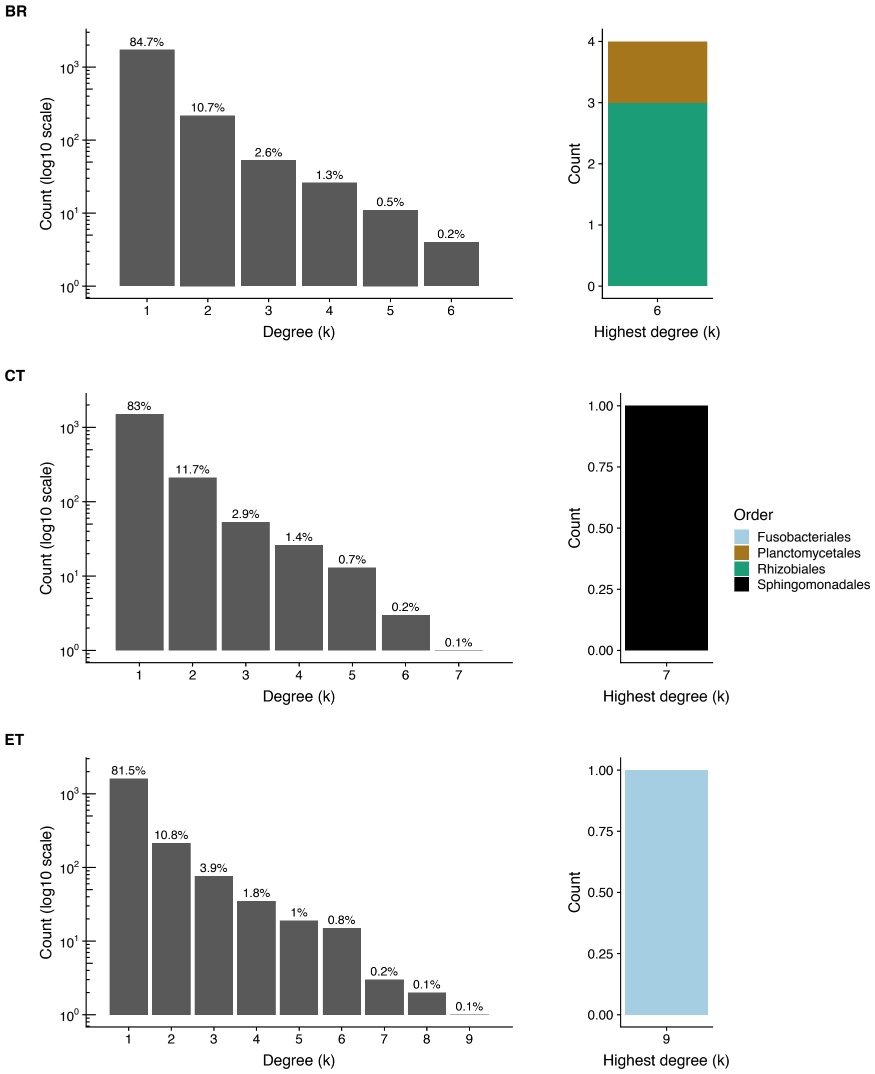


Figure S15. Number of ASVs from sediment for each degree (k) on the left and Order of ASVs present in the highest degree on the right. Each degree represents one water body. ASVs found in only one water body are represented in the category 1. Likewise, ASVs present in sediment across all water bodies are represented in the highest category of each histogram. Note that the count for each histogram is in log10 scale. The Order of each ASV present in sediment across all water bodies from each locality is shown on the barplot on the right. Same color across barplots represent the same Order. BR = Brazil, CT = Central Texas, ET = East Texas.

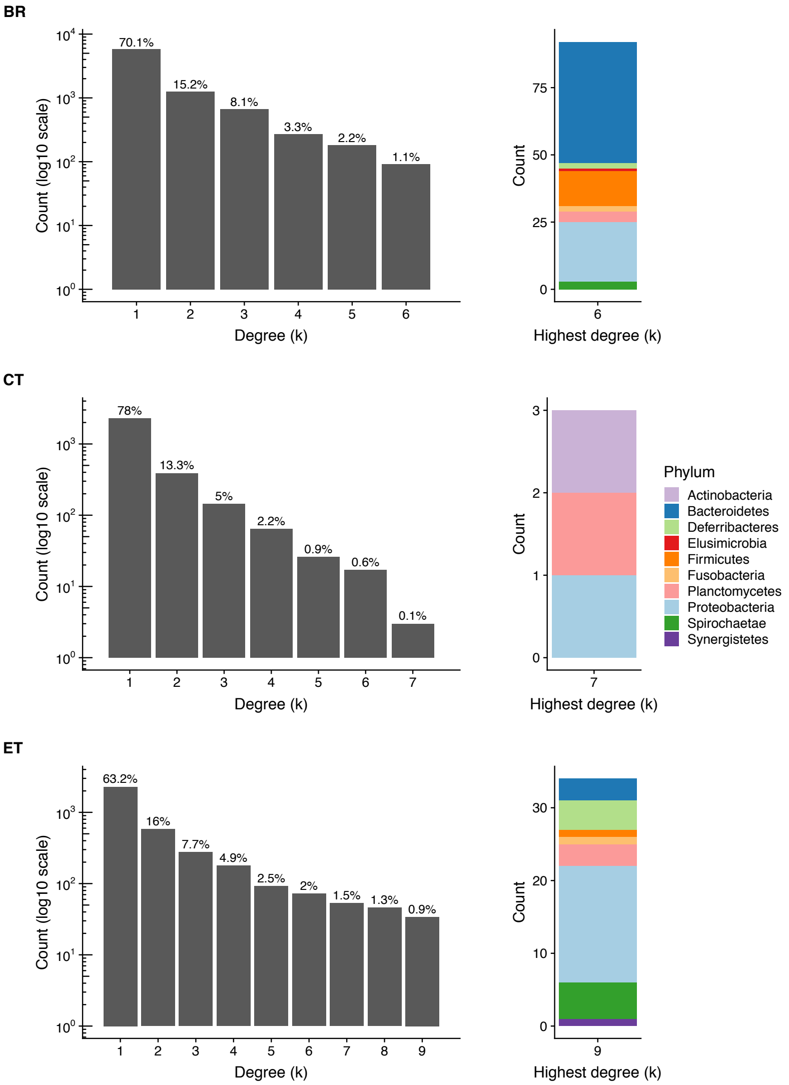


Figure S16. Number of ASVs from tadpoles for each degree (k) on the left and Phylum of ASVs present in the highest degree on the right. Each degree represents one water body. ASVs found in only one water body are represented in the category 1. Likewise, ASVs present in tadpoles across all water bodies are represented in the highest category of each histogram. Note that the count for each histogram is in log10 scale. The Phylum of each ASV present in tadpoles across all water bodies from each locality is shown on the barplot on the right. Same color across barplots represent the same Phylum. BR = Brazil, CT = Central Texas, ET = East Texas.

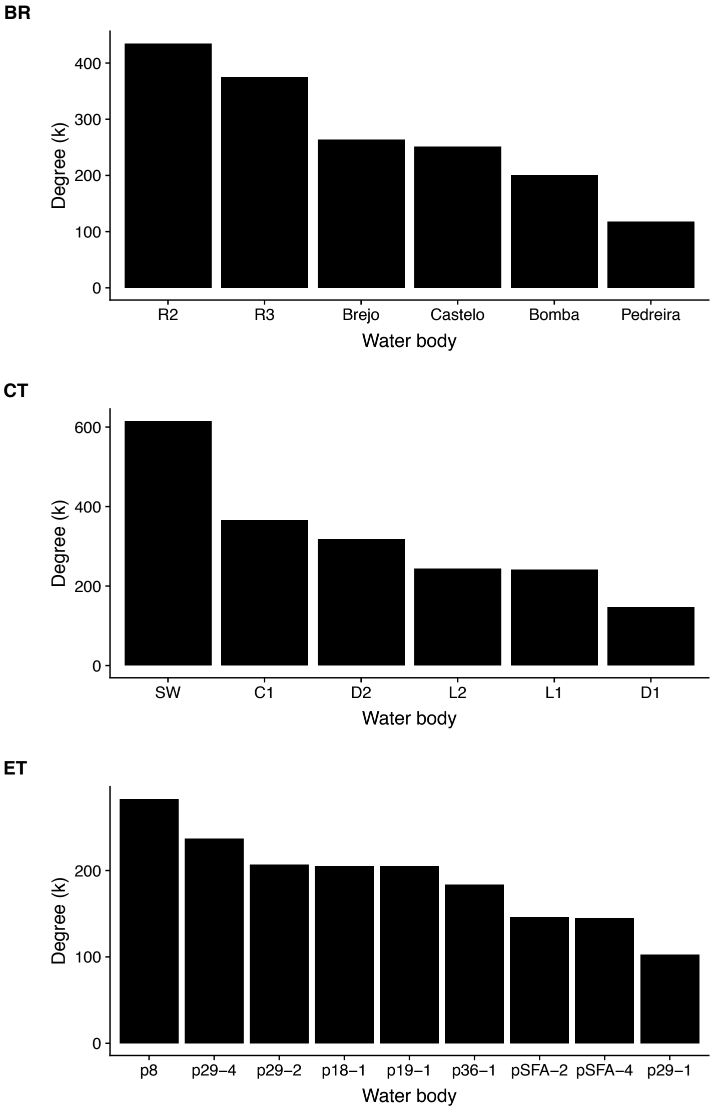


Figure S17. Histograms of number of interactions (degree, k) with ASVs from water per water body. The highest the degree, the more ASVs in that water body.

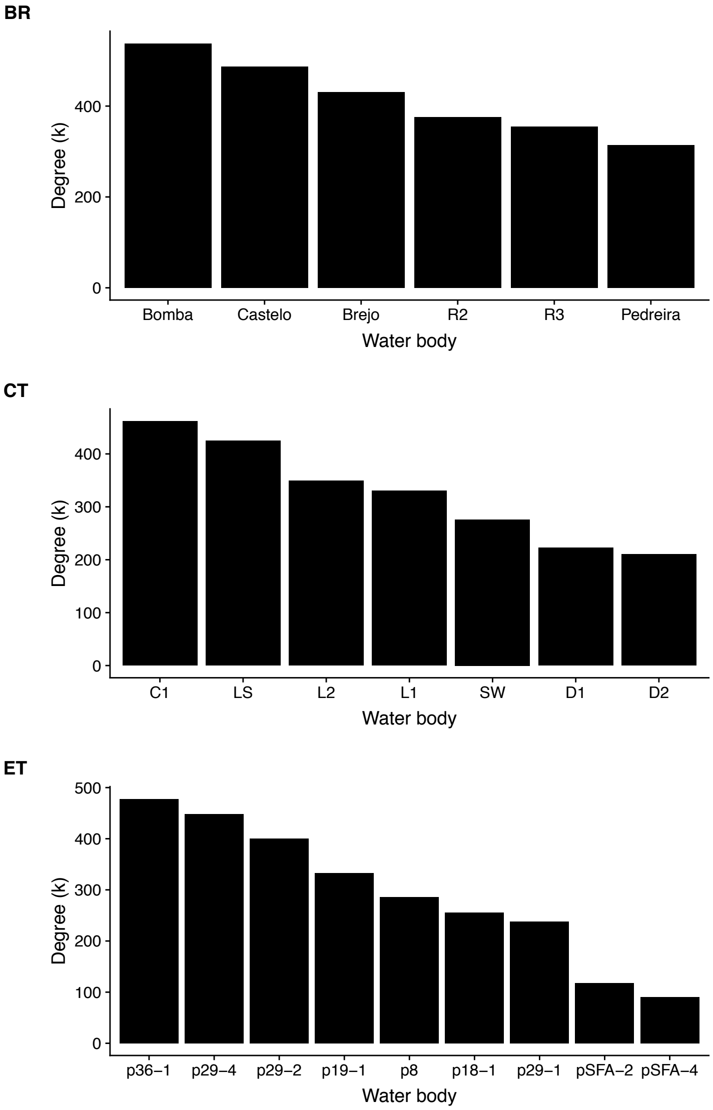


Figure S18. Histograms of number of interactions (degree, k) with ASVs from sediment per water body. The highest the degree, the more ASVs in that water body.

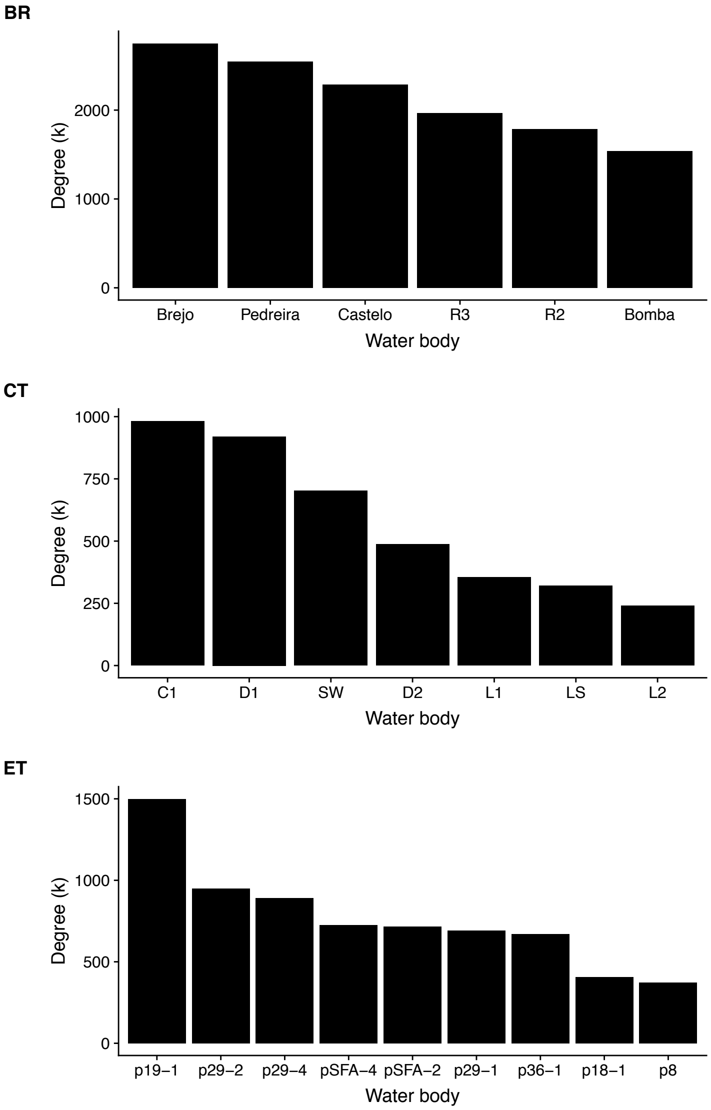


Figure S19. Histograms of number of interactions (degree, k) with ASVs from tadpoles per water body. The highest the degree, the more ASVs in that water body.

**Tables**

Table S1. Species of tadpoles sampled in Brazil (BR), Central (CT) and East Texas (ET) with abbreviations used in figures and text.

| Species | Abbreviations | Location |
|---|---|---|
| *Acris crepitans* | A. crepitans; A. cre | CT, ET |
| *Dendropsophus microps* | D. microps; D. mic | BR |
| *Dendropsophus minutus* | D. minutus; D. min | BR |
| *Gastrophryne carolinensis* | G. carolinensis; G. car | ET |
| *Gastrophryne olivacea* | G. olivacea; G. oli | CT |
| *Hyla versicolor* | H. versicolor; H. ver | ET |
| *Hylodes phyllodes* | H. phyllodes; H. phy | BR |
| *Hylodes* sp. | Hylodes sp.; H. sp. | BR |
| *Hypsiboas faber* | H. faber; H. fab | BR |
| *Hypsiboas pardalis* | H. pardalis; H. par | BR |
| *Phyllomedusa olfersii* | P. olfersii; P. olf | BR |
| *Proceratophrys boiei* | P. boiei; P. boi | BR |
| *Rana berlandieri* | R. berlandieri; R. ber | CT |
| *Rana catesbeiana* | R. catesbeiana; R. cat | ET |
| *Rana clamitans* | R. clamitans; R. cla | ET |
| *Rana sphenocephala* | R. sphenocephala; R. sph | ET |
| *Rhinella* sp. | Rhinella sp.; R. sp | BR |
| *Scinax hayii* | S. hayii; S. hay | BR |
| *Scinax obtriangulatus* | S. obtriangulatus; S. obt | BR |

Table S2. Results of a Permutational Analysis of Variance (perMANOVA) testing differences in composition between samples (999 permutations). Sample type refers to the perMANOVA using all samples (tadpoles, sediment, and water). If a difference was detected we conducted pairwise comparisons with Bonferroni correction (adj. p). The microbiome composition data used is based on Amplicon Sequence Variants (ASVs). BR = Brazil, CT = Central Texas, ET = East Texas.

| ASV | | PHILR | | | Coda | | |
|---|---|---|---|---|---|---|---|
| | | $R^2$ | F | p/ajd. p | $R^2$ | F | p/ajd. p |
| BR | Sample type$_{(2,92)}$ | 0.19 | 10.67 | 0.001 | 0.05 | 2.43 | 0.001 |
| | Tadpole-water$_{(1,86)}$ | 0.1 | 9.23 | 0.003 | 0.02 | 2.07 | 0.015 |
| | Tadpole-sediment$_{(1,86)}$ | 0.13 | 13.14 | 0.003 | 0.03 | 2.92 | 0.003 |
| | Water-sediment$_{(1,11)}$ | 0.09 | 1.01 | 1.000 | 0.12 | 1.31 | 0.027 |
| CT | Sample type$_{(2,38)}$ | 0.14 | 2.88 | 0.001 | 0.11 | 2.12 | 0.001 |
| | Tadpole-water$_{(1,31)}$ | 0.09 | 2.78 | 0.033 | 0.07 | 2.16 | 0.024 |
| | Tadpole-sediment$_{(1,32)}$ | 0.10 | 3.27 | 0.003 | 0.07 | 2.31 | 0.009 |
| | Water-sediment$_{(1,12)}$ | 0.16 | 2.09 | 0.006 | 0.13 | 1.59 | 0.006 |
| ET | Sample type$_{(2,94)}$ | 0.36 | 26.02 | 0.001 | 0.12 | 5.96 | 0.001 |
| | Tadpole-water$_{(1,85)}$ | 0.25 | 27.98 | 0.003 | 0.08 | 7.27 | 0.003 |
| | Tadpole-sediment$_{(1,85)}$ | 0.25 | 27.95 | 0.003 | 0.06 | 5.27 | 0.003 |
| | Water-sediment$_{(1,17)}$ | 0.18 | 3.4 | 0.006 | 0.18 | 3.55 | 0.003 |

Table S3. Results of a Permutational Analysis of Variance (perMANOVA) testing differences in composition between samples (999 permutations). Sample type refers to the perMANOVA using all samples (tadpoles, sediment, and water). If a difference was detected we conducted pairwise comparisons with Bonferroni correction (adj. p). The microbiome composition data used is based on Amplicon Sequence Variants (ASVs) clustered by Genus. BR = Brazil, CT = Central Texas, ET = East Texas.

| Genus | | PHILR | | | Coda | | |
|---|---|---|---|---|---|---|---|
| | | $R^2$ | F | p/ajd. p | $R^2$ | F | p/ajd. p |
| BR | Sample type$_{(2,92)}$ | 0.27 | 16.19 | 0.001 | 0.20 | 11.20 | 0.001 |
| | Tadpole-water$_{(1,86)}$ | 0.14 | 13.70 | 0.003 | 0.11 | 10.59 | 0.003 |
| | Tadpole-sediment$_{(1,86)}$ | 0.19 | 20.43 | 0.003 | 0.14 | 13.60 | 0.003 |
| | Water-sediment$_{(1,11)}$ | 0.19 | 2.41 | 0.027 | 0.20 | 2.41 | 0.003 |
| CT | Sample type$_{(2,38)}$ | 0.17 | 3.78 | 0.001 | 0.22 | 4.91 | 0.001 |
| | Tadpole-water$_{(1,31)}$ | 0.11 | 3.75 | 0.003 | 0.15 | 5.16 | 0.003 |
| | Tadpole-sediment$_{(1,32)}$ | 0.12 | 4.12 | 0.006 | 0.15 | 5.35 | 0.003 |
| | Water-sediment$_{(1,12)}$ | 0.21 | 2.96 | 0.012 | 0.23 | 3.35 | 0.009 |
| ET | Sample type$_{(2,94)}$ | 0.34 | 23.19 | 0.001 | 0.32 | 21.33 | 0.001 |
| | Tadpole-water$_{(1,85)}$ | 0.25 | 27.47 | 0.003 | 0.25 | 27.37 | 0.003 |
| | Tadpole-sediment$_{(1,85)}$ | 0.21 | 21.79 | 0.003 | 0.19 | 19.65 | 0.003 |
| | Water-sediment$_{(1,17)}$ | 0.29 | 6.56 | 0.003 | 0.33 | 7.98 | 0.003 |

Table S4. Results of a Permutational Analysis of Variance (perMANOVA) testing differences in composition between samples (999 permutations). Sample type refers to the perMANOVA using all samples (tadpoles, sediment, and water). If a difference was detected we conducted pairwise comparisons with Bonferroni correction (adj. p). The microbiome composition data used is based on Amplicon Sequence Variants (ASVs) clustered by Family. BR = Brazil, CT = Central Texas, ET = East Texas.

| Family | | PHILR | | | Coda | | |
|---|---|---|---|---|---|---|---|
| | | $R^2$ | F | p/ajd. p | $R^2$ | F | p/ajd. p |
| BR | Sample type$_{(2,92)}$ | 0.21 | 12.05 | 0.001 | 0.25 | 15.02 | 0.001 |
| | Tadpole-water$_{(1,86)}$ | 0.11 | 10.64 | 0.003 | 0.14 | 13.36 | 0.003 |
| | Tadpole-sediment$_{(1,86)}$ | 0.16 | 15.85 | 0.003 | 0.18 | 18.95 | 0.003 |
| | Water-sediment$_{(1,11)}$ | 0.14 | 1.60 | 0.513 | 0.25 | 3.27 | 0.003 |
| CT | Sample type$_{(2,38)}$ | 0.27 | 6.49 | 1.000 | 0.28 | 7.02 | 0.001 |
| | Tadpole-water$_{(1,31)}$ | | | | 0.19 | 6.94 | 0.003 |
| | Tadpole-sediment$_{(1,32)}$ | | | | 0.21 | 8.27 | 0.003 |
| | Water-sediment$_{(1,12)}$ | | | | 0.28 | 4.20 | 0.009 |
| ET | Sample type$_{(2,94)}$ | 0.38 | 27.6 | 0.001 | 0.37 | 26.61 | 0.001 |
| | Tadpole-water$_{(1,85)}$ | 0.27 | 30.84 | 0.003 | 0.28 | 32.87 | 0.003 |
| | Tadpole-sediment$_{(1,85)}$ | 0.26 | 29.01 | 0.003 | 0.24 | 26.53 | 0.003 |
| | Water-sediment$_{(1,17)}$ | 0.22 | 4.62 | 0.003 | 0.35 | 8.50 | 0.003 |

Table S5. Results of a Permutational Analysis of Variance (perMANOVA) testing if the identity of the species (local effects) of the identity of the pond (regional effects) are related to the gut microbiome composition of tadpoles grouping Amplicon Sequence Variants (ASVs) by their Genus. Models were fitted to CODA and PHILR transformed data. The significance of the marginal effects was tested based on 999 permutations. BR = Brazil, CT = Central Texas, ET = East Texas.

| Genus | | PHILR | | | Coda | | |
|---|---|---|---|---|---|---|---|
| | | $R^2$ | F | p | $R^2$ | F | p |
| BR | Water body$_{(4,80)}$ | 0.09 | 2.7 | 0.001 | 0.12 | 3.6 | 0.001 |
| | Species$_{(9,80)}$ | 0.19 | 2.5 | 0.001 | 0.17 | 2.3 | 0.001 |
| CT | Water body$_{(5,25)}$ | 0.53 | 7.9 | 0.001 | 0.49 | 5.3 | 0.001 |
| | Species$_{(1,25)}$ | 0.05 | 3.5 | 0.001 | 0.09 | 2.6 | 0.015 |
| ET | Water body$_{(8,76)}$ | 0.29 | 5.1 | 0.001 | 0.26 | 4.0 | 0.001 |
| | Species$_{(5,76)}$ | 0.18 | 5.1 | 0.001 | 0.17 | 4.2 | 0.001 |

Table S6. Results of a Permutational Analysis of Variance (perMANOVA) testing if the identity of the species (local effects) of the identity of the pond (regional effects) are related to the gut microbiome composition of tadpoles grouping Amplicon Sequence Variants (ASVs) by their Family. Models were fitted to CODA and PHILR transformed data. The significance of the marginal effects was tested based on 999 permutations. BR = Brazil, CT = Central Texas, ET = East Texas.

| Family | | PHILR | | | Coda | | |
|---|---|---|---|---|---|---|---|
| | | $R^2$ | F | p | $R^2$ | F | p |
| BR | Water body$_{(4,80)}$ | 0.09 | 2.5 | 0.006 | 13.6 | 4.3 | 0.001 |
| | Species$_{(9,80)}$ | 0.19 | 2.3 | 0.001 | 15.7 | 2.2 | 0.001 |
| CT | Water body$_{(5,25)}$ | 0.57 | 9.55 | 0.001 | 0.49 | 5.4 | 0.001 |
| | Species$_{(1,25)}$ | 0.07 | 3.72 | 0.002 | 0.05 | 2.7 | 0.009 |
| ET | Water body$_{(8,76)}$ | 0.25 | 4.7 | 0.001 | 0.29 | 4.7 | 0.001 |
| | Species$_{(5,76)}$ | 0.22 | 6.6 | 0.001 | 0.16 | 4.2 | 0.001 |

Table S7. Results of a variation partitioning analysis testing the unique local (species identity) or regional (pond identity) effects as well their shared effects on the tadpole gut microbiome grouping the Amplicon Sequence Variants (ASVs) by their Genus. Significance test base on 1000 permutations, except for the shared component, which cannot be tested. BR = Brazil, CT = Central Texas, ET = East Texas.

| Genus | | PHILR | | Coda | |
|---|---|---|---|---|---|
| | | Variation explained | p | Variation explained | p |
| | Regional | 0.06 | 0.001 | 0.10 | 0.001 |
| BR | Local | 0.12 | 0.001 | 0.10 | 0.001 |
| | Regional + local | 0.16 | not testable | 0.14 | not testable |
| | Regional | 0.51 | 0.001 | 0.43 | 0.001 |
| CT | Local | 0.04 | 0.001 | 0.04 | 0.005 |
| | Regional + local | 0.11 | not testable | 0.07 | not testable |
| | Regional | 0.25 | 0.001 | 0.21 | 0.001 |
| ET | Local | 0.16 | 0.001 | 0.14 | 0.001 |
| | Regional + local | 0.05 | not testable | 0.04 | not testable |

Table S8. Results of a variation partitioning analysis testing the unique local (species identity) or regional (pond identity) effects as well their shared effects on the tadpole gut microbiome grouping the Amplicon Sequence Variants (ASVs) by their Family. Significance test base on 1000 permutations, except for the shared component, which cannot be tested. BR = Brazil, CT = Central Texas, ET = East Texas.

| Family | | PHILR | | Coda | |
|---|---|---|---|---|---|
| | | Variation explained | p | Variation explained | p |
| | Regional | 0.06 | 0.007 | 0.12 | 0.001 |
| BR | Local | 0.11 | 0.005 | 0.09 | 0.001 |
| | Regional + local | 0.12 | not testable | 0.16 | not testable |
| | Regional | 0.55 | 0.001 | 0.43 | 0.001 |
| CT | Local | 0.11 | 0.002 | 0.04 | 0.005 |
| | Regional + local | 0.04 | not testable | 0.07 | not testable |
| | Regional | 0.21 | 0.001 | 0.25 | 0.001 |
| ET | Local | 0.21 | 0.001 | 0.14 | 0.001 |
| | Regional + local | 0.07 | not testable | 0.03 | not testable |

Table S9. Results of multiple regression on distance matrices testing the correlation between microbiome samples (PHILR and CODA transformed) from water or sediment and spatial distance between water bodies, taking into account a distance matrix based on environmental variables (BR and ET only; see main text for details). BR = Brazil, CT = Central Texas, ET = East Texas.

| | | BR | | | | CT | | | | ET | | | |
|---|---|---|---|---|---|---|---|---|---|---|---|---|---|
| | | PHILR | | CODA | | PHILR | | CODA | | PHILR | | CODA | |
| | | Value | p | Value | p | Value | p | Value | p | Value | p | Value | p |
| Water | Environmental | 4.9 | 0.041 | 3.9 | 0.004 | na | na | na | na | 8.4 | 0.006 | -2.4 | 0.726 |
| | Spatial | -71 | 0.517 | -167.4 | 0.236 | 11.9 | 0.004 | 73.9 | 0.003 | -1.6 | 0.739 | -3 | 0.730 |
| | R2 | 0.22 | 0.145 | 0.2 | 0.15 | 0.33 | 0.004 | 0.34 | 0.003 | 0.28 | 0.029 | 0.01 | 0.889 |
| | F | 1.78 | 0.145 | 1.6 | 0.15 | 6.4 | 0.004 | 6.57 | 0.003 | 6.3 | 0.029 | 0.2 | 0.889 |
| Sediment | Environmental | 0.23 | 0.413 | 1.7 | 0.065 | na | na | na | na | 4.8 | 0.009 | 11.6 | 0.336 |
| | Spatial | 9.3 | 0.471 | 20.8 | 0.924 | 5.7 | 0.062 | -0.59 | 0.979 | 0.8 | 0.666 | -17.3 | 0.304 |
| | R2 | 0.09 | 0.407 | 0.04 | 0.718 | 0.19 | 0.062 | 6.28 | 0.979 | 0.27 | 0.044 | 0.12 | 0.357 |
| | F | 0.66 | 0.407 | 0.22 | 0.718 | 4.44 | 0.062 | 0.001 | 0.979 | 6.23 | 0.044 | 2.42 | 0.357 |

Table S10. Modules assigned to each water body based on the algorithm FastGreedy. Metanetworks are water bodies connected by shared bacteria (amplicon sequence variant - ASV) found in the gut of tadpoles, water, or sediment. BR = Brazil, CT = Central Texas, ET = East Texas.

| | Water body | Tadpole | Water | Sediment |
|---|---|---|---|---|
| BR | Bomba | 1 | 0 | 4 |
| | Brejo | 4 | 1 | 3 |
| | Castelo | 3 | 2 | 5 |
| | Pedreira | 5 | 4 | 0 |
| | R2 | 2 | 5 | 2 |
| | R3 | 0 | 3 | 1 |
| CT | C1 | 3 | 0 | 6 |
| | D1 | 2 | 1 | 0 |
| | D2 | 1 | 2 | 2 |
| | L1 | 4 | 3 | 4 |
| | L2 | 4 | 3 | 3 |
| | LS | 4 | NA | 5 |
| | SW | 0 | 4 | 1 |
| ET | 18-1 | 6 | 4 | 2 |
| | 19-1 | 8 | 6 | 3 |
| | 29-1 | 1 | 0 | 1 |
| | 29-2 | 3 | 1 | 5 |
| | 29-4 | 2 | 4 | 8 |
| | 36-1 | 7 | 5 | 7 |
| | 8 | 0 | 7 | 0 |
| | SFA-2 | 5 | 3 | 4 |
| | SFA-4 | 4 | 2 | 6 |
